# Cdc14 activates autophagy to regulate spindle pole body dynamics during meiosis

**DOI:** 10.1101/2021.07.21.453268

**Authors:** Wenzhi Feng, Orlando Argüello-Miranda, Suhong Qian, Fei Wang

**Affiliations:** UT Southwestern Medical Center, Department of Cell Biology, Dallas, TX, 75390, USA; UT Southwestern Medical Center, Lyda Hill Department of Bioinformatics, Dallas, TX, 75390, USA

**Keywords:** autophagy, meiosis, sporulation, phosphatase, Cdc14, Atg13, Atg1, Spo74, SPB, cell development

## Abstract

Autophagy, a conserved eukaryotic lysosomal degradation pathway that responds to environmental and cellular cues, is regulated by multiple signaling pathways that oversee cell survival, growth, and proliferation. In budding yeast, autophagy plays an essential role in meiotic exit, although the molecular mechanisms underlying its regulation and cargo selection remain unknown. In this study, we found that autophagy is maintained during meiosis and is upregulated at anaphase I and anaphase II. In addition, we found that cells with higher levels of autophagy during meiosis I and II completed meiosis faster, and that genetically activated autophagy machinery increased sporulation efficiency. Strikingly, our data revealed that Cdc14, a highly conserved phosphatase that counteracts Cdc28 (CDK1), is a meiosis-specific autophagy regulator. At anaphase I and anaphase II, Cdc14 was activated and released from the nucleolus into the cytoplasm, where it dephosphorylated Atg13 to stimulate Atg1 kinase activity and thus autophagy. Importantly, the meiosis-specific spindle pole body (SPB, the yeast centrosome) component (Spo74) was sensitized to autophagy-mediated degradation at anaphase II, upon its dephosphorylation by Cdc14. Together, our findings reveal a meiosis-tailored mechanism of Cdc14 that spatiotemporally guides meiotic autophagy activity to control SPB dynamics.

## Introduction

Meiosis depends on a carefully orchestrated sequence of intracellular processes that produce viable gametes after a diploid cell undergoes two rounds of nuclear division. Among several cellular processes specific to meiosis, targeted proteolysis provides a rapid and irreversible mechanism for promoting transitions between meiotic stages (Neiman, 2011, King et al., 1996). Autophagy is a highly conserved lysosomal degradation pathway that was originally discovered as a response to cellular stress. Although autophagy is best characterized as a survival mechanism, emerging evidence indicates that this versatile degradation mechanism is also involved in selective regulation of cell development and differentiation (Politi et al., 2014, Rojansky et al., 2016, Sato and Sato, 2011, Wang et al., 2020b). In budding yeast, an important model system for research on meiosis, autophagy is required for meiotic entry (Piekarska et al., 2010, Sarkar et al., 2014, Wen et al., 2016, Enyenihi and Saunders, 2003, Straub et al., 1997) and exit (Wang et al., 2020b, Wang et al., 2020a). Specifically, loss of autophagy during meiotic cell division results in delayed onset of meiosis II (metaphase II and anaphase II), failed exit from meiosis, and cell death (Wang et al., 2020b). In the absence of autophagy, the cell over-amplifies the spindle pole body (SPB, the yeast centrosome), the nucleation center for the spindle, and the membrane of the prospore (the precursor of the daughter cell), resulting in abnormal chromosome segregation and prospore membrane formation (Wang et al., 2020b). However, the mechanisms underlying these important roles of autophagy remain elusive. Autophagy at anaphase I rapidly destroys the RNA-binding protein (RBP) Rim4 (Wang et al., 2020b), which forms amyloid-like aggregates to sequester a subset of mRNAs (Berchowitz et al., 2015). Autophagy-dependent removal of Rim4 allows the timely translation of Rim4-sequestered mRNAs, including those encoding key mid-late meiotic regulators (e.g., the B-type cyclin Clb3) and essential players in sporulation (Berchowitz et al., 2013, Berchowitz et al., 2015). Thus, autophagy may influence SPB dynamics via Rim4-controlled translation (Wang et al., 2020b). To date, however, Rim4 is the only protein identified as an autophagy substrate specific to meiosis. It remains unknown whether autophagy can directly control SPB number during meiosis, thus safeguarding proper chromosome segregation and sporulation, by degrading SPB-related protein(s).

The clearance of intracellular Rim4 aggregates at anaphase I and the apparent role of autophagy in regulating SPB dynamics at meiosis exit (anaphase II) imply that autophagic activity at these two meiotic stages is important and should be secured by meiosis. What, then, is the mechanism? As a major degradation pathway, autophagy must be tightly regulated to avoid excess or insufficient autophagic degradation. Consistent with this, dysregulation of autophagy is associated with a wide variety of diseases and disorders (Mizushima and Komatsu, 2011, Meijer and Codogno, 2009, Guo et al., 2013). Various signal transduction mechanisms have been implicated in the regulation of autophagy in response to extra-and intracellular stimuli. One important and highly conserved strategy for regulating autophagy regulation is posttranslational modification (PTM). In particular, alteration of protein phosphorylation state can rapidly switch autophagy initiation on and off (Funakoshi et al., 1997, Scott et al., 2000, Kamada et al., 2000, Memisoglu et al., 2019, Mao et al., 2013, Pengo et al., 2017, Davis et al., 2016, Licheva et al., 2021) as well as influence cargo selection (Aoki et al., 2011, Pfaffenwimmer et al., 2014, Gatica et al., 2018, Stolz et al., 2014). To date, several protein kinases have been identified and mechanistically characterized in regulating autophagy, including mTOR complex 1 (mTORC1), a master suppressor of autophagy initiation (Kim and Guan, 2015). In sharp contrast, our knowledge of the role of protein phosphatases in autophagy regulation remains much more limited(Ohsumi, 2005).

Intriguingly, in mammalian mitosis, autophagy is regulated by the mitotic master kinase CDK1 (Cdc28 in yeast), which replaces the role of mTORC1 in phosphorylating autophagy initiation factors such as ULK1 (Atg1 in yeast), ATG13 (Atg13 in yeast), and ATG14 (Atg14 in yeast) (Odle et al., 2020). Whether CDK1 activates or suppresses mitotic autophagy during mitosis remains controversial (Yamasaki et al., 2020, Li et al., 2020). Similarly, we do not yet know whether CDK1 modulates autophagy during meiosis, a process in which programmed gene expression, protein dynamics, and homeostasis differ from those in mitosis. Nonetheless, TORC1 activity is suppressed during meiosis to allow meiotic entry and progression (Yamamoto, 2004, van Werven and Amon, 2011, Harigaya and Yamamoto, 2007). The main phosphatase that counteracts CDK1 is Cdc14. CDK1 and Cdc14, which are activated in metaphase and anaphase, respectively, are positioned to shape autophagy activity in coordination with meiotic progression. Surprisingly, inactivation of *CDC14* in haploid vegetative yeast cells is associated with a reduction in autophagy after TORC1 inactivation (starvation), a condition that is not directly related to mitosis or meiosis (Kondo et al., 2018). It is unclear whether Cdc14 directly regulates autophagy under this condition; it is essential in regulating numerous signaling pathways (Breitkreutz et al., 2010), but its known activities are almost exclusively related to the cell cycle and cell division (Odle et al., 2020, Yamasaki et al., 2020, Li et al., 2020). Hence, this finding implies that during meiotic anaphase, autophagy could be regulated by Cdc14.

By antagonizing CDK1 activity during mitosis and meiosis, the members of the Cdc14 family of dual phosphatases regulate the duplication cycle of SPBs in yeast and centrosomes in humans (Wu et al., 2008, Avena et al., 2014, Elserafy et al., 2014). In HeLa human cervical cancer cells and BJ and MRC-5 human fibroblasts, hCdc14B depletion by RNA interference causes abnormal centriole amplification, whereas overexpression of hCdc14B prevents unscheduled duplication of centriole in cells arrested in prolonged S phase (Wu et al., 2008). During the transition from meiosis I to meiosis II, Cdc14 is involved in re-licensing SPBs (Fox et al., 2017). The underlying mechanism of these observations remains largely unclear, although it is known that in *S. cerevisiae*, Cdc14 dephosphorylates Sfi1, a half-bridge complex protein, to periodically license the SPB duplication during mitosis (Avena et al., 2014, Elserafy et al., 2014). Among the most striking phenotypes of autophagy inhibition during meiotic cell divisions is abnormal amplification of SPBs, raising the possibility that Cdc14 coordinates with autophagy to control meiotic SPB dynamics.

In this study, we found that Cdc14 stimulates autophagy in anaphase I and anaphase II. During both anaphase I and anaphase II, Cdc14 relocates from the nucleolus into the cytoplasm, where it dephosphorylates Atg13 to activate Atg1 and thus autophagy. We identified two motifs of Atg13 that are required for Cdc14–Atg13 binding and six serine residues of Atg13 that are key targets of Cdc14 phosphatase activity, providing molecular insights into the Cdc14–Atg13 interaction. Importantly, we found that Spo74, a meiosis-specific SPB component, was dephosphorylated by Cdc14 after mid-late anaphase II, making it more sensitive to autophagy-mediated degradation. Together, our findings reveal a meiosis-specific mechanism of autophagy regulation by Cdc14 that directs autophagy to modulate meiotic SPBs by priming Spo74’s autophagic degradation.

## Results

### Autophagy is upregulated in anaphase I and II

Several studies have implied that autophagy-mediated proteolysis is downregulated during meiosis I and II (Piekarska et al., 2010, Sarkar et al., 2014, Wen et al., 2016, Enyenihi and Saunders, 2003, Straub et al., 1997) and replaced by the ubiquitin-proteasome system (UPS) (Wen et al., 2016). However, we have observed that autophagy during meiosis I and II remains essential for meiotic progression and exit (Wang et al., 2020b). To investigate how autophagy is regulated during meiosis, we started by monitoring autophagy flux from meiotic entry to exit and early sporulation in the budding yeast strain W303. We employed a GFP-Atg8 processing assay that quantifies the delivery of GFP-Atg8 on the inner autophagosomal membrane to the vacuole. Atg8 is degraded in the vacuole lumen, leaving behind free GFP, which is more resistant to hydrolysis (Klionsky et al., 2016). We induced GFP-Atg8 (*pZD: GFP-Atg8*) expression by treating cells with β-estradiol at meiotic entry (Fig. 1A) and analyzed GFP-Atg8 processing over a time course by immunoblotting (IB: α-GFP) (Fig. S1A; 1B, top panel). Meiotic stages were determined by immunofluorescence (IF) analysis of spindle microtubules (Fig. 1B, bottom panel) and by IB to detect Ndt80 (Fig. 1B, top panel; S1B), a transcription factor for mid-late meiotic genes expressed shortly before entry into metaphase I. We found that autophagy flux, measured as the GFP-Atg8 processing rate, was relatively stable through early meiotic stages until the onset of meiotic divisions (meiosis I and II) (Fig. 1B). Intriguingly, autophagy during meiotic divisions changed dramatically over time, as reflected by free GFP levels (Fig. 1B, top panel, orange line), suggesting that autophagy might be regulated according to the specific needs of each meiotic stage.

**Figure 1.**
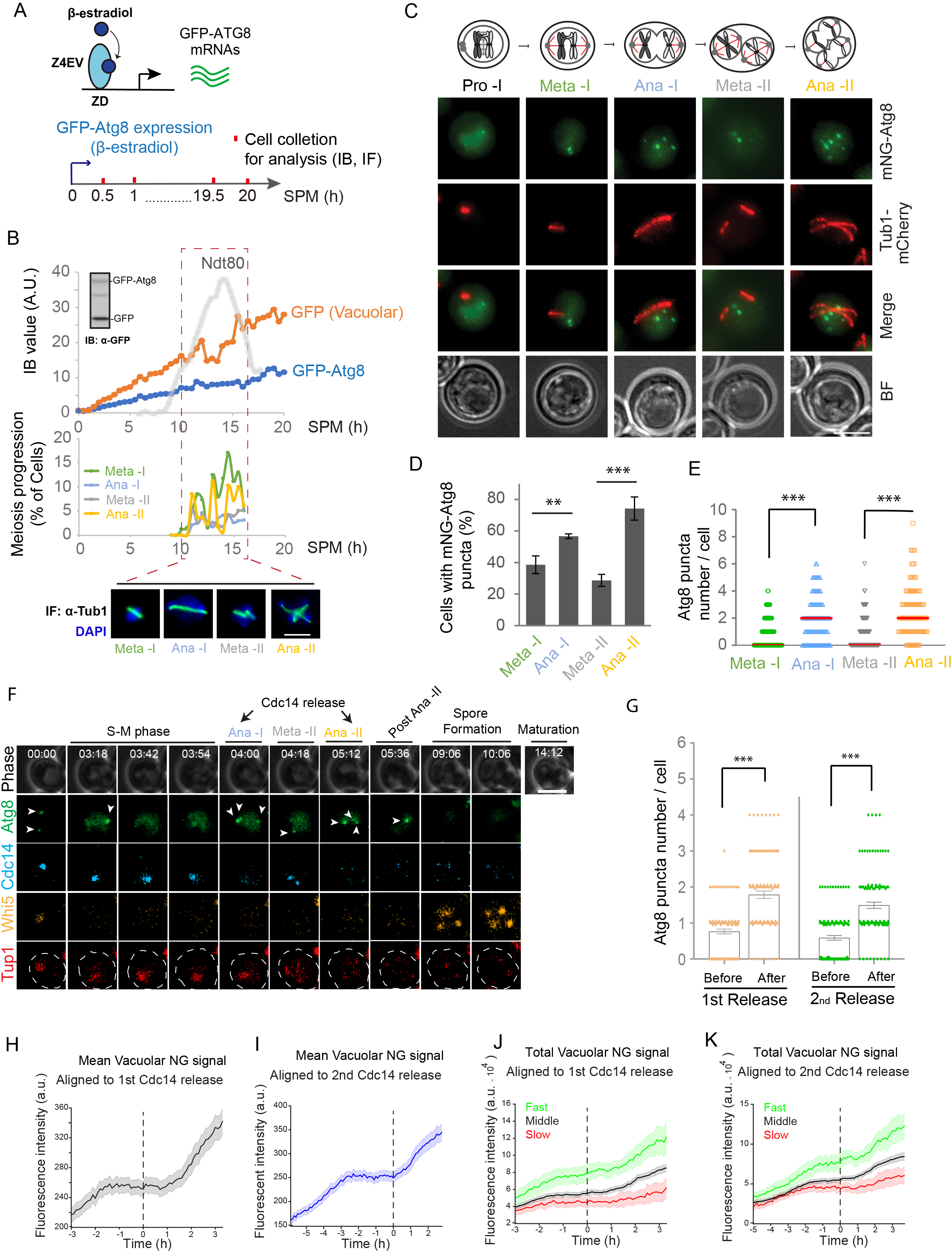
Autophagy is upregulated at anaphase I and II. (A) Schematic of GFP-Atg8 induction and cell collection for immunoblotting (IB) and immunofluorescence staining (IF: α-Tubulin) during meiosis and sporulation. Z4EV is a β-estradiol activated transcription factor and ZD is a Z4EV-driven promoter. (B) Top, graph of cellular GFP-Atg8 (and free GFP) and Ndt80 protein levels (determined by IB, S1A & S1B) during meiosis and sporulation as (A). Bottom, representative images and graph of meiosis stages determined by IF (α-Tubulin). Nuclear DNA was stained by DAPI. Dashed box (red), the time window of meiotic divisions. >100 cells from two independent experiments were analyzed. Scale bar: 5µm. (C-E) Fluorescence microscopy (FM) analysis of mNG-Atg8 puncta and Tub1-mCherry in live cells during Gal-NDT80–synchronized meiosis. The mNG-Atg8 and Tub1-mCherry were expressed under their endogenous promoters. Meiotic stages were identified by the specific Tub1-mCherry distribution pattern depicted on top. (C) Representative images showing composite of five planes (Z-step size, 0.5µm). Scale bar, 5μm. (D) & (E) Quantitation of (C). n ≥ 300 cells at each stage from 3 replicate experiments. Herein, the asterisk represents statistically significant differences: *, p≤0.05; **, p ≤0.01, ***, p≤ 0.001, while error bars represent standard deviations (SD) of 3 replicate experiments. Red line in (E) indicates the median. (F-K) Time-lapse FM of a cell undergoing meiosis. Cell expresses Whi5-mKOk, mNG-Atg8, Cdc14-mTFP1, and Tup1-mNeptune2.5. (F) Representative images. Time, h:min. Scale Bar: 5μm. (G-H) Quantitation of (F) showing mNG-Atg8 puncta number per cell before and after Cdc14 cytosolic releases (G) (n =100 cells from two independent experiments; p ≤0.001, paired t-test; error bar, SEM), or mNG-Atg8 vacuolar signal (line) with the standard deviations (shade) aligned to indicated Cdc14 releases (H-K). In (J & K), cells from (F) were divided into three groups based on how fast they went through meiosis (Green, 10% of the total, the fastest; Pink, 10% of the total, the slowest; Grey, 80% of the total, the middle), (Total, n= 302 cells; KS test).

We then examined autophagy in each stage of meiotic cell division. Given that the time of metaphase I entry differs by ∼1–2 h among individual cells (Nachman et al., 2007), we adapted an inducible *GAL-NDT80* system (Fig. S1C) that enables synchronized metaphase I entry (Fig. S1D) (Carlile and Amon, 2008, Benjamin et al., 2003, Wang et al., 2020b). Using fluorescence microscopy (FM), we monitored endogenously expressed mNeonGreen-Atg8 (mNG-Atg8), which forms puncta, as a quantifiable marker of autophagosome biogenesis (Klionsky et al., 2016). In parallel, we determined meiotic stage based on the characteristic morphology of spindle microtubules labeled with Tub1-mCherry (Fig. 1C). As shown in Fig. 1D, the percentage of cells with mNG-Atg8 puncta was significantly higher in anaphase I and II (57 ± 1.5% and 74 ± 7.4%, respectively) than in metaphase I and II (39 ± 5.6% and 29 ± 3.8%, respectively). Furthermore, the number of mNG-Atg8 puncta per cell was also significantly higher in anaphase I and II (Fig. 1E), indicating that autophagosome biogenesis is upregulated at anaphase I and II.

To track autophagosome biogenesis and autophagic flux through meiosis in single cells, we performed time-lapse FM in a strain endogenously expressing mNG-Atg8 as an autophagy marker. In addition, we tracked Tup1-mNeptune2.5, a nuclear marker; Whi5-mKOκ, a Cdk1 activity marker that is sequestered in the cytoplasm during meiotic divisions but translocates to the nucleus during pre-meiotic G1 and meiotic exit; and Cdc14-mTFP1, whose cytosolic relocation from the nucleolus marks anaphase I and II (Fig. 1F and Movie S1) (Argüello-Miranda et al., 2018). We tracked 302 cells (n=4) that successfully progressed through meiosis and sporulation over the course of 30 h. The number of mNG-Atg8 puncta per cell significantly increased in anaphase I and II, as indicated by two sequential releases of nuclear Cdc14-mTFP1 into the cytosol (Fig. 1F and 1G), providing further confirmation that autophagosome biogenesis increases in anaphase I and II.

To determine the dynamics of autophagosomes, we measured the vacuolar mNG signal (autophagy flux indicator) that was aligned to anaphase I (first Cdc14 release) or anaphase II (second Cdc14 release), using a computational algorithm to track the vacuolar compartment on phase-contrast images (details in Experimental Procedures) (Fig. 1H and 1I). This analysis revealed that the greatest increase in autophagy flux started at anaphase II (Fig. 1I). mNG-Atg8 puncta formation increased at anaphase I (Fig. 1D, 1E, and 1G), whereas autophagy flux (vacuolar mNG signal) did not accelerate until anaphase II (Fig. 1I). This delay likely occurred because autophagosome biogenesis is only briefly upregulated at anaphase I, but suppressed again at the following metaphase II (Fig. 1D and 1E).

Remarkably, the fastest cells to complete meiosis (upper 10% of cells according to time of meiotic completion) also had significantly (p<0.05, KS-Test) higher levels of autophagy flux during meiotic cell divisions (Fig. 1J and 1K, green line). Conversely, the slowest cells to complete meiosis (lower 10% of cells according to time of meiotic completion) also had significantly (p<0.05, KS-Test) lower levels of autophagy flux during meiotic cell divisions (Fig. 1J & 1K, red line). Collectively, these observations indicate that autophagy levels during meiotic cell divisions correlate with the time required to complete meiosis. Moreover, induced expression of a gain-of-function mutant of Atg13 (Atg13-8SA) (Kamada et al., 2010) during meiotic cell divisions upregulated autophagy (Fig. S1E), leading to increased sporulation efficiency (Fig. S1F), whereas inhibition of autophagy by 1NM-PP1 (Blethrow et al., 2004, Wang et al., 2020b) (details in Fig. S2A) abolished sporulation (Fig. S1F), indicating that autophagy levels during meiotic divisions influence meiosis and sporulation. Together, our data demonstrate that autophagy is dynamically regulated during meiotic divisions, with autophagosome biogenesis peaking at anaphase I and II. Furthermore, autophagy levels during meiotic cell divisions correlate with the efficiency of meiosis progression.

### Atg1 activity is stimulated in anaphase I and II

The increase in autophagosome biogenesis in anaphase I and II led us to analyze the autophagy master kinase Atg1, whose activation is required for autophagy initiation. Inhibition of Atg1 kinase activity during meiosis suppressed autophagy and inhibited sporulation (Fig. S1F) (Wang et al., 2020b). Thus, Atg1 was a strong candidate for the factor responsible for upregulating autophagy in anaphase I and II. Hence, we examined Atg1-as kinase activity during GAL-NDT80– synchronized meiotic divisions, specifically asking whether Atg1 activity is upregulated at anaphase I and II. We induced synchronized meiotic divisions in cells expressing a functional analog–sensitive allele of Atg1, Atg1-M102G (Atg1-as) (Kamber et al., 2015); the mutant protein has an enlarged ATP binding pocket that can accommodate a bulky ATP γS analog (6-PhEt-ATP-γ-S) as a substrate to specifically phosphorylate Atg1 substrates, including itself (auto-phosphorylation), in cell lysates derived from meiotic cell divisions (Fig. 2A). When we monitored Atg1-as auto-phosphorylation, we found that Atg1 kinase activity periodically increased, peaking at ∼2, ∼4, and ∼6 h after Ndt80 induction (Fig. 2B), whereas Atg1 protein levels remained stable (Fig. S2B and S2C).

**Figure 2.**
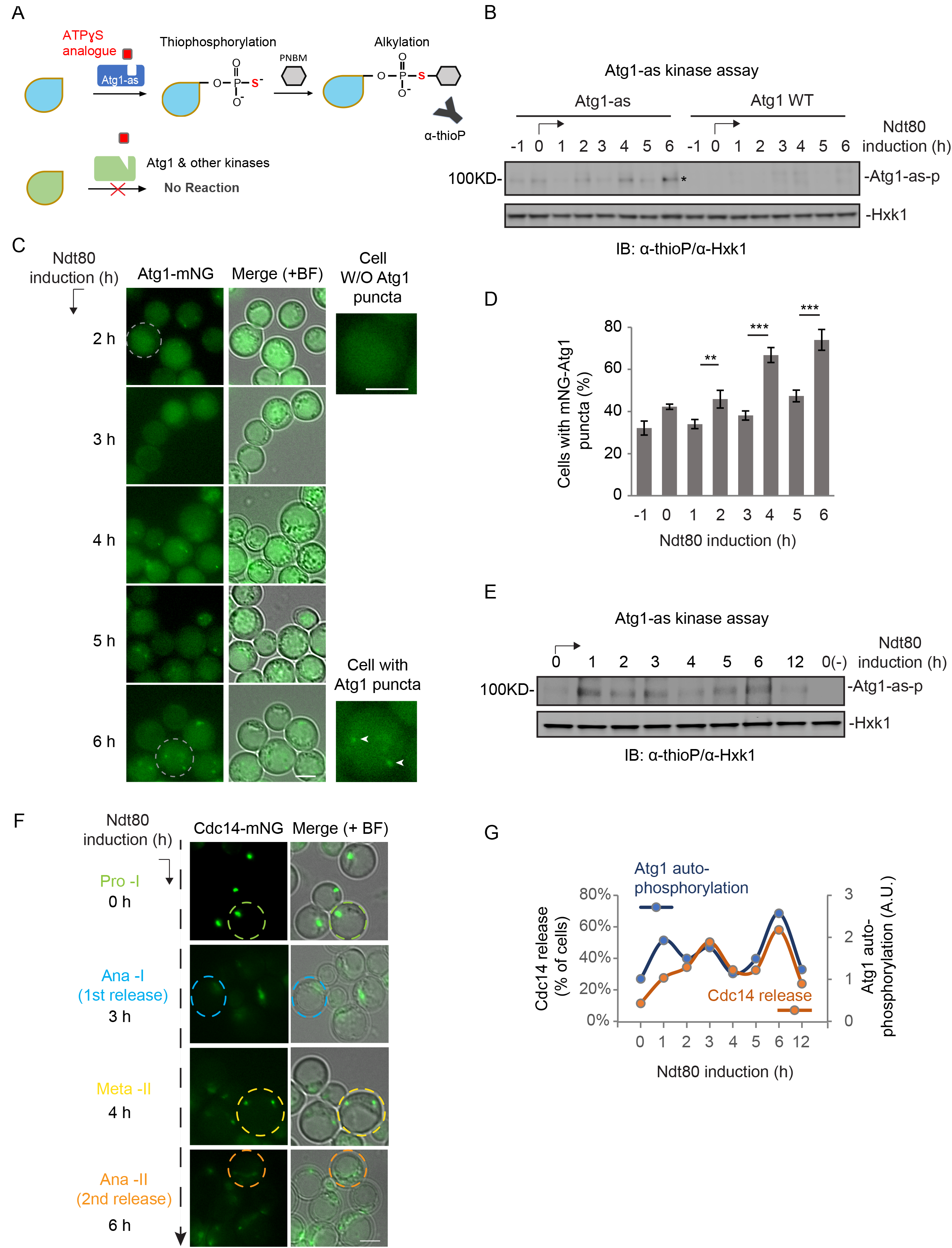
Atg1 activity is stimulated in anaphase I and II. (A) Schematic of the chemical-genetic strategy for monitoring Atg1-as (analog-sensitive Atg1, Atg1-M102G) kinase activity *in vitro*. Atg1-as thiophosphorylates its substrates with a bulky ATPγS analog (N6-PhEt-ATP-γ-S). Thiophosphorylated substrates of Atg1-as can then be alkylated with para-nitrobenzyl mesylate (PNBM) and detected by IB using anti-thiophosphate ester (α-thioP) antibodies. (B) Atg1-as or Atg1 cells during Gal-Ndt80–synchronized meiosis were treated as in (A). Following Atg1 kinase assay, the whole-cell lysates were subjected to immunoblotting (IB) with indicated antibodies. Unless indicated otherwise, 1μM β-estradiol was added to meiotic cells after 12 h in SPM to synchronize the entry of meiotic divisions, depicted as t=0 h of Ndt80 induction. Hxk1, hexokinase isoenzyme 1 (loading control); *, Atg1-as auto-phosphorylation. (C & D) Synchronized meiotic cells expressing Atg1-mNG were analyzed by FM. Representative FM Images (C) and graph showing the percent of cells with detectable Atg1-mNG puncta (D) (n ≥ 300 cells at each time point from 3 replicate experiments; t-test). Representative Cells with/without Atg1-mNG puncta, circled by grey dashed lines, are shown on the right. Scale bars, 5μm. (E & F) Cells expressing Cdc14-mNG were subjected to IB analysis for Atg1-as auto-phosphorylation (E) and FM analysis for Cdc14-mNG cytosolic relocation (F). (E) The experiment was performed as in (A). Minus (-), alkylation in the absence of ATP. (F) Cells circled by dashed lines are representative of indicated meiotic stages. Scale bar, 5μm. (G) Quantitation of (E & F). Graphs show Atg1-as autophosphorylation (blue line) and Cdc14-mNG cytosolic release (orange, n ≥ 300).

Next, we investigated whether elevated Atg1 activation during meiotic divisions leads to increased autophagy initiation. Upon activation, Atg1 is recruited to the pre-autophagosomal structure (PAS), marked by cytosolic puncta of Atg1 (Yamamoto et al., 2016). Therefore, we performed FM to examine Atg1-mNG puncta formation during GAL-NDT80–synchronized meiosis. Consistent with the pattern of Atg1 auto-phosphorylation (Fig. 2B), the proportion of cells with Atg1-mNG puncta increased at ∼2, ∼4, and ∼6 h after Ndt80 induction (Fig. 2C and 2D). Thus, Atg1 was periodically stimulated and recruited to PAS. Next, we asked whether Atg1 stimulation correlates with anaphase I and II progression. We measured Atg1 kinase activity in meiotic cell lysates by IB (Fig. 2E; 2G, blue line), and determined anaphases based on the relocation of nuclear Cdc14-mNG into the cytosol (Fig. 2F; 2G, orange line). Remarkably, this experiment revealed that Atg1 kinase activity and the cytosolic release of Cdc14 co-fluctuated (Fig. 2G), indicating that anaphase I and II progression is coupled with an increase in Atg1 activity.

### Cdc14 stimulates Atg1 activity in anaphase to assist autophagy initiation

The tight spatiotemporal correlation between Atg1 activity and Cdc14 raised the possibility that Cdc14 directly activates Atg1 in the cytosol by dephosphorylating Atg13, the main regulatory component of the Atg1 complex. If so, Cdc14 should bind or be close to Atg13 or the Atg1–Atg13 complex. Therefore, we performed FM analysis to investigate whether Cdc14-mNG colocalizes with cytosolic mScarlet-Atg13 puncta during meiotic divisions at the PAS, where Atg1 is activated. Even though Cdc14 is only transiently in contact with its substrates (Rudolph, 2007, Wang et al., 2004), we found that 13±2% of cells containing mScarlet-Atg13 puncta showed mScarlet-Atg13/ Cdc14-mNG colocalization (Fig. 3A and 3B). Consistent with this, Cdc14-mNG also colocalized with Atg1-mScarlet puncta with similar efficiency (Fig. 3C and 3D, 12±1%). Importantly, colocalization did not occur during prophase I even after prolonged prophase arrest, indicating that anaphase-triggered cytosolic presence of Cdc14 is required for Cdc14/Atg13-Atg1 colocalization at PAS (Fig. 3A and 3B; 3C and 3D). Thus, during anaphase, Cdc14 is close to Atg1 and Atg13 at PAS and is consequently appropriately localized to dephosphorylate Atg13 to activate Atg1.

**Figure 3.**
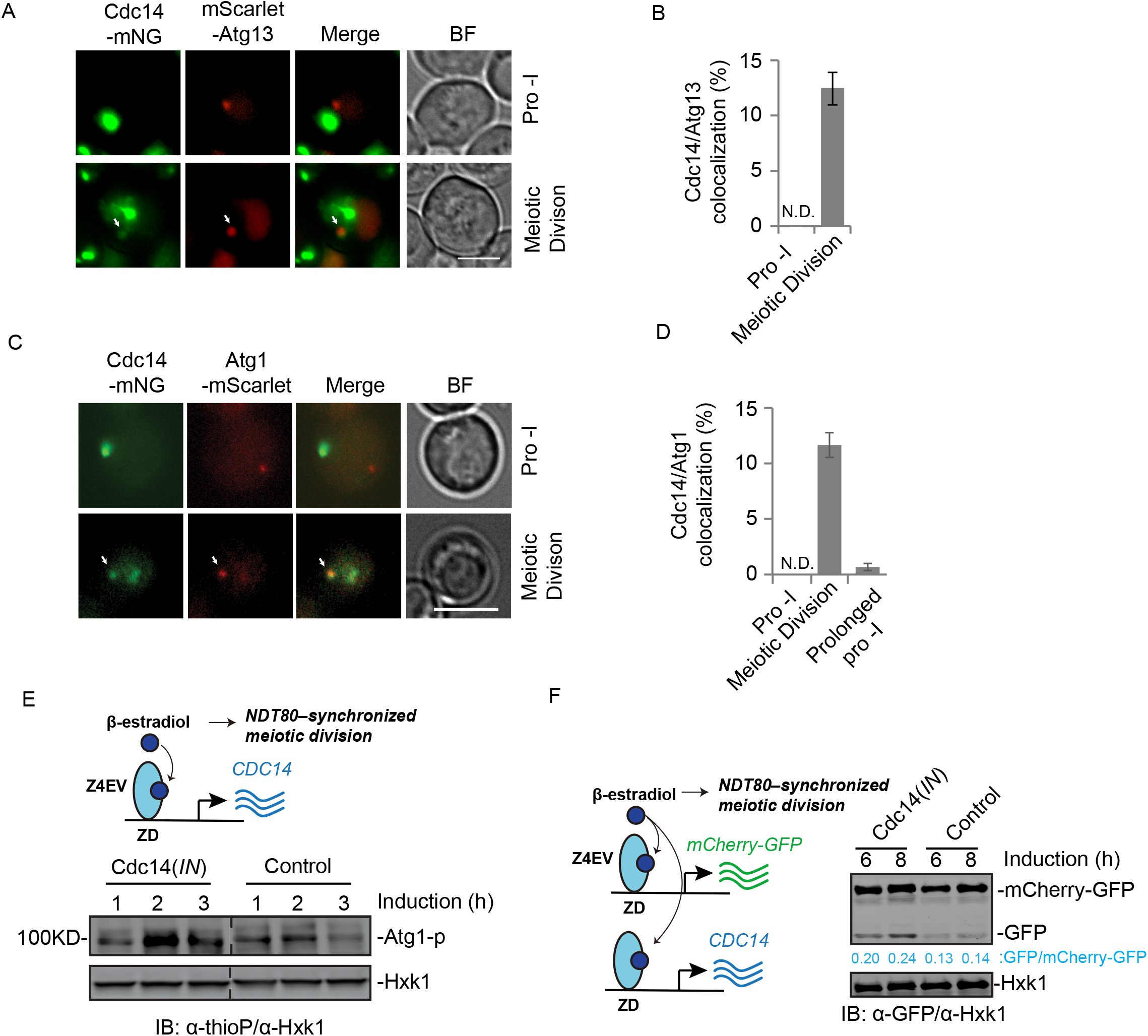
Cdc14 stimulates Atg1 activity in anaphase to assist autophagy initiation. (A-D) FM analysis of cells expressing indicated genes under their endogenous promoters. Shown are representative FM images of Gal-Ndt80–synchronized meiotic cells at prophase-I (Before Ndt80 induction) and during meiotic divisions (After Ndt80 induction). Cdc14-mNG/Atg1-mScarlet, or Cdc14-mNG/mScarlet-Atg13 colocalization is indicated by white arrows in (A & C) and quantified in (B & D). Scale bars, 5μm. Cells that carry Atg1-mScarlet (C) or mScarlet-Atg13 (D) puncta were analyzed (n≥300 cells from 3 replicate experiments, t-test). (E) Atg1-as activity in response to Cdc14(*IN*) was examined as in part (2A & 2B). Top, schematic of induced Cdc14(*IN*) expression (pZD: Cdc14) during synchronized meiosis. (F) IB analysis with indicated antibodies, showing the change of mCherry-GFP cleavage level during meiotic divisions upon induced Cdc14(*IN*). Left, schematic of simultaneously induced expression of Cdc14, mCherry-GFP and Ndt80.

To determine whether Atg1 is activated by Cdc14, we induced expression of extra Cdc14 (*pZD: CDC14*), herein referred to as Cdc14(*IN*), to increase cytosolic Cdc14 levels (Bouchoux and Uhlmann, 2011, Higuchi and Uhlmann, 2005, Visintin et al., 1998, Miller et al., 2015) immediately before entry into metaphase I (Fig. S3A & S3B). This strategy ensures WT sporulation efficiency while avoiding the potential negative effects of extreme Cdc14 overexpression during meiosis (Fig. S3C). Cdc14 induction enhanced Cdc14/Atg13 colocalization (Fig. S3D and S3E, 35±5%; as opposed to 13 ± 2% in Fig. 3B); stimulated Atg1 activity, as judged by Atg1 kinase assay (Fig. 3E), and increased vacuolar processing of GFP-Atg8 (Fig. S3F) and mCherry-GFP (Fig. 3F), a non-selective bulk autophagy reporter. These results suggest that during meiotic cell divisions, Cdc14 modulates autophagy by activating Atg1, a key event in autophagy initiation.

To determine whether Cdc14 can stimulate autophagy independent of meiosis, we investigated Cdc14/Atg13 colocalization under conditions unrelated to meiosis. When Cdc14-mNG and Atg13-mScarlet were expressed under the control of the corresponding endogenous promoters, we observed no colocalization of puncta during proliferation (log-phase) or when autophagy was stimulated by stress stimuli such as nitrogen starvation (Fig. S3G), indicating that Cdc14 activity might specifically modulates autophagy during meiotic divisions.

### Cdc14 dephosphorylates Atg13 to activate Atg1

To investigate the possibility that Cdc14 activates Atg1 to initiate autophagy, we examined the effect of recombinant Cdc14 on Atg1 kinase activity *in vitro*. First, we confirmed that recombinant Cdc14 is active using a colorimetric phosphatase assay (Fig. S4A and S4B). 6His-3FLAG-Cdc14 exhibited concentration-and time-dependent phosphatase activity (Fig. S4C) that was inhibited by phosphatase inhibitors (PI) (Fig. S4D). Cdc14m (6His-3FLAG-Cdc14m) did not exhibit detectable phosphatase activity even at 80 μg/ml (1.12 µM) (Fig. S4E). Next, we immunoisolated FLAG-Atg1 complex from yeast cell lysates (Fig. S4F) and treated it with recombinant Cdc14. Each cell contains ∼8550 molecules of Cdc14 (∼0.4µM) (Ghaemmaghami et al., 2003), which remains cytosolic for ∼20 min during meiotic divisions. In a dose-dependent manner ([Cdc14] ranging from 0.3 to 0.7 µM), Atg1 auto-phosphorylation (Atg1 activity) increased in response to 20 min of Cdc14 phosphatase activity (Fig. 4A). In addition, phosphatase inhibitor fully abolished the effect of Cdc14 on Atg1 auto-phosphorylation (Fig. 4A). This observation suggests that Cdc14 can directly dephosphorylate Atg1-associated protein(s) in order to activate Atg1.

**Figure 4.**
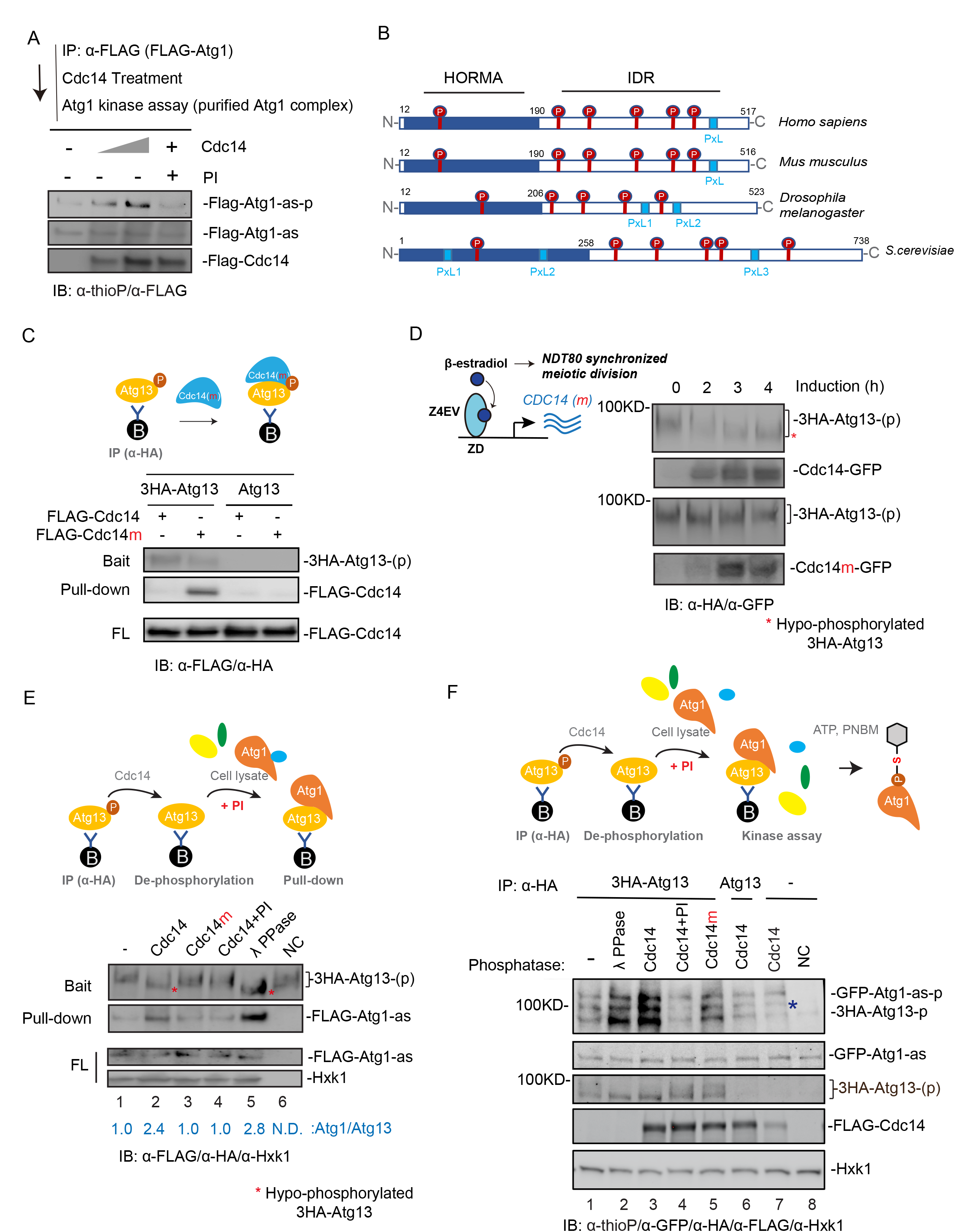
Cdc14 dephosphorylates Atg13 to activate Atg1. (A) FLAG-Atg1-as protein was affinity-purified (IP: α-FLAG) from extracts of meiotic cells arrested at Prophase I, and then treated as indicated, followed by Atg1-as kinase assay as diagrammed (2A). PI, Phosphatase Inhibitors. (B) Protein domain organization of Atg13 from various species, with the location of protein domains annotated by residue number. Shown are the Serine-Proline (SP, in red) and Proline-x-Leucine (PxL, light blue box) sites. IDR, Intrinsically <u>D</u>isordered <u>R</u>egions; HORMA, for <u>Ho</u>p1p, <u>R</u>ev7p and <u>MA</u>D2. (C) Top, schematic of experimental design. Extracts from cells with indicated genetic backgrounds under vegetative growth were subjected to immunoprecipitation (IP: α-HA). Shown are binding of recombinant FLAG-Cdc14 or FLAG-Cdc14m to bead-immobilized 3HA-Atg13, overnight at 4 °C. FL, flow through. (D) Top, schematic of experimental design. IB analysis of 3HA-Atg13-p dephosphorylation (band shift) after Cdc14-GFP(*IN*) or Cdc14m-GFP(*IN*) was induced during synchronized meiotic divisions. 3HA-Atg13-p/3HA-Atg13 in cell lysates were enriched by IP (α-HA), followed by IB (α-HA). Left, schematic of induced Cdc14-GFP(*IN*)/Cdc14m-GFP(*IN*) expression. *, hypo-phosphorylated 3HA-Atg13. (E & F) Top, schematic of experimental design. Bead-immobilized 3HA-Atg13-p from IP (α-HA) was treated by Cdc14 or as indicated to enable dephosphorylation, followed by incubation with *Flag-Atg1-as* cell lysate to pull down Flag-Atg1-as (E) or with *GFP-Atg1-as* cell lysates to stimulate Atg1 auto-phosphorylation (F). Samples were resolved by SDS-PAGE followed by IB with indicated antibodies. (E) NC, no cell lysate. (F) NC, minus ATP. *, unknown identity.

In the Atg1 kinase complex, dephosphorylation of Atg13 generally enhances the Atg13–Atg1 interaction and activates Atg1 at PAS (Kamada et al., 2000, Memisoglu et al., 2019, Yeasmin et al., 2016). We inspected the amino acid sequence of *S. cerevisiae* Atg13 and observed several Cdc14 preferred dephosphorylation sites (i.e., SP sites) (Gray and Barford, 2003, Bremmer et al., 2012) and putative Cdc14 docking sites (PxL sites) (Kataria et al., 2018) (Fig. 4B). Therefore, we investigated whether Cdc14 binds to and dephosphorylates Atg13. Due to the transient nature of the Cdc14/substrate interaction, it is difficult to be detected in co-immunoprecipitation (Co-IP) assays (Flint et al., 1997, Bradshaw and Dennis, 2009). To circumvent this limitation, we employed the catalytically dead mutant Cdc14m, which stabilizes Cdc14 binding to its *bona fide* substrates (Bloom et al., 2011, Chen et al., 2013), and performed IP of 3HA-Atg13 to pull down supplemented recombinant Cdc14 (6His-3FLAG-Cdc14) or Cdc14m (6His-3FLAG-Cdc14m). At the same concentration, Cdc14m, but not Cdc14, was pulled down by 3HA-Atg13 (Fig. 4C), supporting the idea that Atg13 binds to Cdc14 and could be a Cdc14 substrate. Consistent with this, induction of Cdc14(*IN*) at metaphase I entry increased the mobility of the 3HA-Atg13 band on the gel (IB: α-HA) (Fig. 4D), whereas Cdc14m(*IN)* had no effect, suggesting that 3HA-Atg13 is dephosphorylated by Cdc14 during meiotic divisions.

To directly probe whether Atg13 is a substrate of Cdc14 phosphatase, we reconstituted Cdc14-mediated Atg13 dephosphorylation *in vitro*. We isolated hyper-phosphorylated Atg13 (3HA-Atg13-p) by IP from lysates of vegetative cells grown in SD, and then treated with either Cdc14, Cdc14m, or λ phosphatase in the presence or absence of phosphatase inhibitors (PI). Cdc14 treatment accelerated the gel migration of the band corresponding to 3HA-Atg13-p on the gel to a similar degree as λ phosphatase treatment (Fig. 4E, lane 2 and 5, red asterisk). By contrast, neither Cdc14m nor Cdc14 with phosphatase inhibitors (PI) detectably dephosphorylated 3HA-Atg13-p (Figure 4E, lane 3 & 4).

Next, we investigated whether Atg13 dephosphorylation by Cdc14 activates Atg13, and hence Atg1. 3HA-Atg13 dephosphorylated by Cdc14 treatment exhibited enhanced binding to Atg1 in cell lysates (Figure 4E, compare lanes 1 & 2). Furthermore, the dephosphorylated 3HA-Atg13 greatly stimulated the kinase activity of Atg1 (GFP-Atg1-as) in vegetative cell lysates (YPD medium) (Figure 4F, lane 1 & 3) in a manner dependent on Cdc14 phosphatase activity (Figure 4F, lane 2, 4 &5). Together, these results demonstrate that Cdc14 dephosphorylates Atg13, enhancing the Atg13/Atg1 interaction and Atg1 activation.

### Two PxL motifs in Atg13 are required for Cdc14 docking

The primary sequence of Atg13 contains three PxL sites that are potentially accessible to Cdc14 docking (Fig. 5A and S5A) (Kataria et al., 2018). We individually mutated each of the PxL sites to AxG (PxL1m, PxL2m, and PxL3m), which disrupts substrate binding of Cdc14 (Kataria et al., 2018). We then immunoprecipitated the Atg13 variants (3HA-tagged wild type and PxL mutants) from cell lysates and probed for binding of supplemented recombinant Cdc14m (6His-3FLAG-Cdc14m) after unbound proteins were washed away. Only WT and PxL2m, but not PxL1m or PxL3m, pulled down Cdc14m, suggesting that the PxL1 and PxL3 sites are required for Cdc14 binding (Fig. 5B).

**Figure 5.**
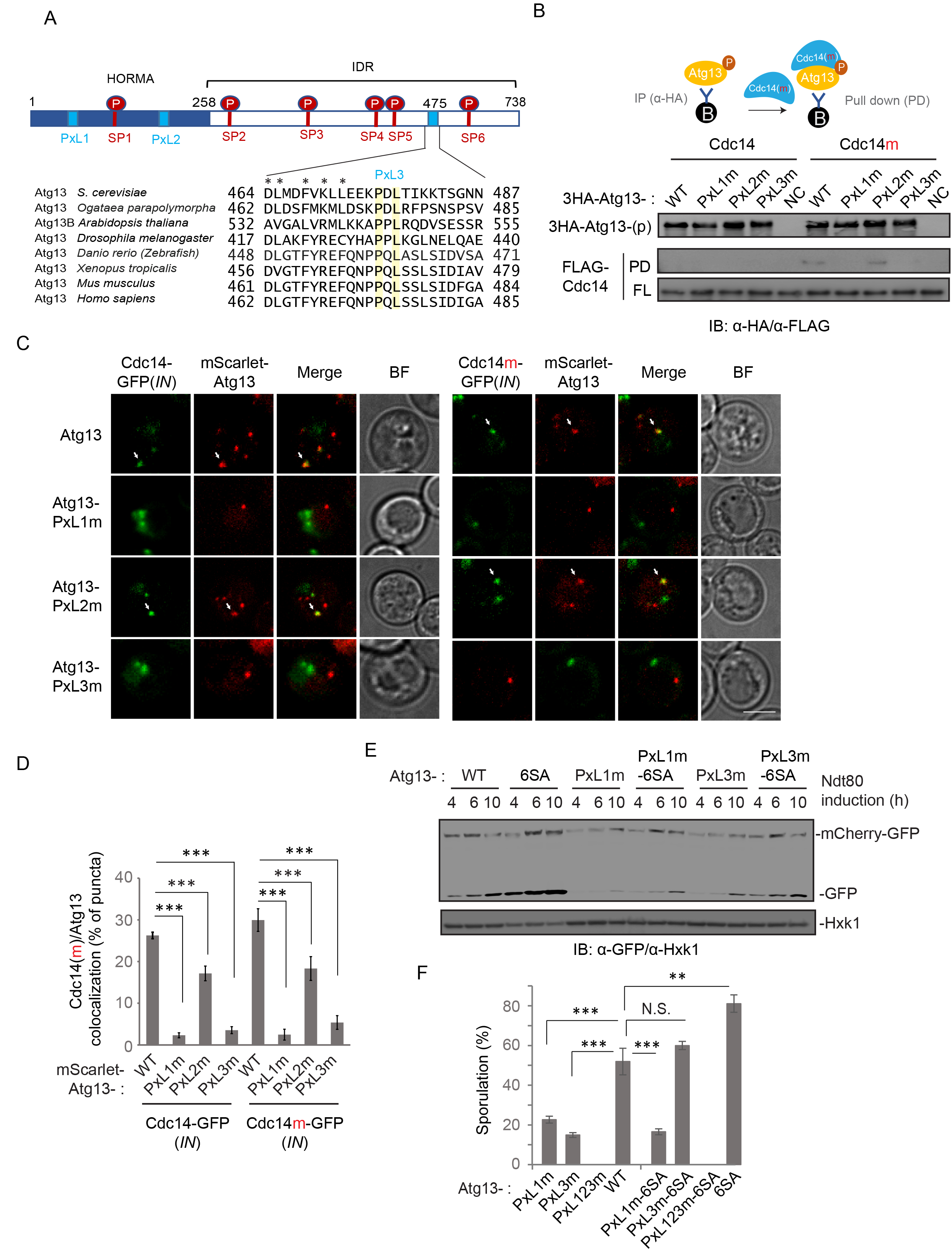
Two PxL motifs in Atg13 are required for Cdc14 docking. (A) Schematic of Atg13 with SP sites and PxL sites. SP sites and PxL sites, in need, are mutated to AP and AxG, respectively. Bottom, amino acid sequence alignment of the PxL3 region of Atg13 homologs from indicated species. Identical residues are highlighted in yellow and highly similar residues are asterisk-labeled. (B) IB analysis of recombinant FLAG-tagged Cdc14/Cdc14m binding to bead-immobilized 3HA-Atg13 variant proteins that were purified by IP (α-HA) from vegetative cell lysates. Top, schematic of experimental design. (C & D) representative FM images (C) and quantitation (D) of GFP-tagged Cdc14(*IN*)/Cdc14m(*IN*) colocalization with mScarlet-tagged Atg13 variants, during meiotic divisions (n≥300 cells, t-test). White arrows indicate colocalization. Scale bar, 5μm. (E) IB of extracts from meiotic cells with indicated antibodies. Cell simultaneously expresses mCherry-GFP (p*ZD: mCherry-GFP*) and Ndt80, induced by 1µM β-estradiol. (F) Shown are the percent of cells of indicated Atg13 variants showing sporulation after 48 h in SPM (n≥300 cells; t-test).

Intriguingly, all three PxL mutants decreased Atg13 puncta formation during meiotic cell divisions (Fig. S5B), indicating a reduction in autophagy initiation. To determine whether the reduced autophagy initiation observed in the PxL mutants could be rescued by stimulating Atg1, we exposed meiotic cells to rapamycin, which stimulates autophagy by inhibiting TORC1 (Kamada et al., 2000). During meiotic divisions, rapamycin only has a moderate effect; however, Cdc14(*IN*) in combination with rapamycin restores Atg13 puncta formation in PxLm cells (Fig. S5B), suggesting that PxLm-induced defects in autophagy initiation are due to a weakened Cdc14– Atg13 interaction. Next, we examined Cdc14–Atg13 puncta colocalization in rapamycin-treated meiotic cells expressing mScarlet-Atg13 and Cdc14-GFP (*IN*) or Cdc14m-GFP (*IN*). Consistent with our pull-down results (Fig. 5B), PxL1m and PxL3m greatly reduced the colocalization of cytosolic Cdc14-GFP (or Cdc14m-GFP) with mScarlet-Atg13 puncta during meiotic divisions, whereas PxL2m had a less pronounced but still significant effect (Fig. 5C and 5D: wild type, 26±0.8%; PxL1m, 2±0.6%; PxL2m, 17±1.8%; PxL3m, 4±0.8%). Collectively, our data demonstrate that the PxL1 and PxL3 sites in Atg13 are required for Cdc14-Atg13 interaction and autophagy initiation; the role of PxL2 in Cdc14–Atg13 interaction is less critical. Consistent with this, Pxl1m and Pxl3m, but not PxL2m, significantly decreased autophagy levels during meiosis (Fig. S5C) and also decreased sporulation efficiency by ∼60% relative to the wild type, and PxL123m (triple mutations) failed to sporulate altogether (Fig. S5D).

Genomic replacement of *ATG13* by the phospho-defective mutants *ATG13-2SA* or *ATG13-6SA* [containing two (2SA) or six (6SA) Serine to Alanine mutations at the SP sites, respectively] is sufficient to stimulate autophagy in cells grown in SD medium, with 6SA having a greater effect (Fig. S5E). Furthermore, 6SA boosted autophagy during meiosis (Fig. S5F). Because the *ATG13-6SA* mutant mimics the product of Cdc14-mediated Atg13 dephosphorylation, we asked whether it could rescue Atg13-PxL1m or Atg13-PxL3m mutants that failed to bind Cdc14 and potentially remained hyper-phosphorylated. Indeed, autophagic degradation of mCherry-GFP (Fig. 5E) and sporulation (Fig. 5F) was restored to the wild-type level in PxL3m-6SA cells, demonstrating that PxL3 is a *bona fide* Cdc14-docking site on Atg13 that enables Atg13 dephosphorylation. Interestingly, PxL1m was not rescued by 6SA. It is possible that Cdc14 docking at the PxL1 site of Atg13 might play a non-enzymatic structural role at the PAS site; alternatively, it may be involved in dephosphorylating other Atg proteins (e.g., Atg1) at the PAS. It is worth noting that PxL3, which resides in an intrinsically disordered region (IDR), is the only PxL site in Atg13 that is conserved across species and likely has the greatest access to the six SP sites due to their physical proximity and flexibility (Fig. 5A).

### Cdc14 primes Spo74 for autophagic degradation at meiotic exit

Given that autophagy inhibition leads to anomalous spindle pole body (SPB) duplication after anaphase II (Wang et al., 2020b), when the greatest upregulation of autophagy flux occurs (Fig. 1I), we investigated whether SPB proteins are anaphase II–specific autophagy substrates. To explore this possibility, we first determined the subcellular localization of Cdc14-mNG, Spo74-mTFP, and mScarlet-Atg13 during meiotic divisions. Spo74, a meiosis-specific SPB protein that resides on the cytosolic face of SPB complexes, is essential for prospore (daughter cell) membrane biogenesis. As shown in Fig. 6A, Cdc14-mNG at anaphase II resided near Spo74 puncta (Fig. 6B, 15±1%) and Atg13 (Fig. 6C, 13±1.8%) during late meiosis. Intriguingly, 1NM-PP1 increased the colocalization of Atg13/Spo74 (Fig. 6D, from 7±1% to 11±2%), reflecting a possible Spo74-PAS(Atg13) interaction that was enhanced when Atg1 was inhibited. Remarkably, in ∼15% of 1NM-PP1–treated cells, extra Spo74 puncta (>4) at the end of meiosis were present, indicating Spo74 accumulation and SPB aberrations in the absence of autophagy (Fig. 6E & 6F).

**Figure 6.**
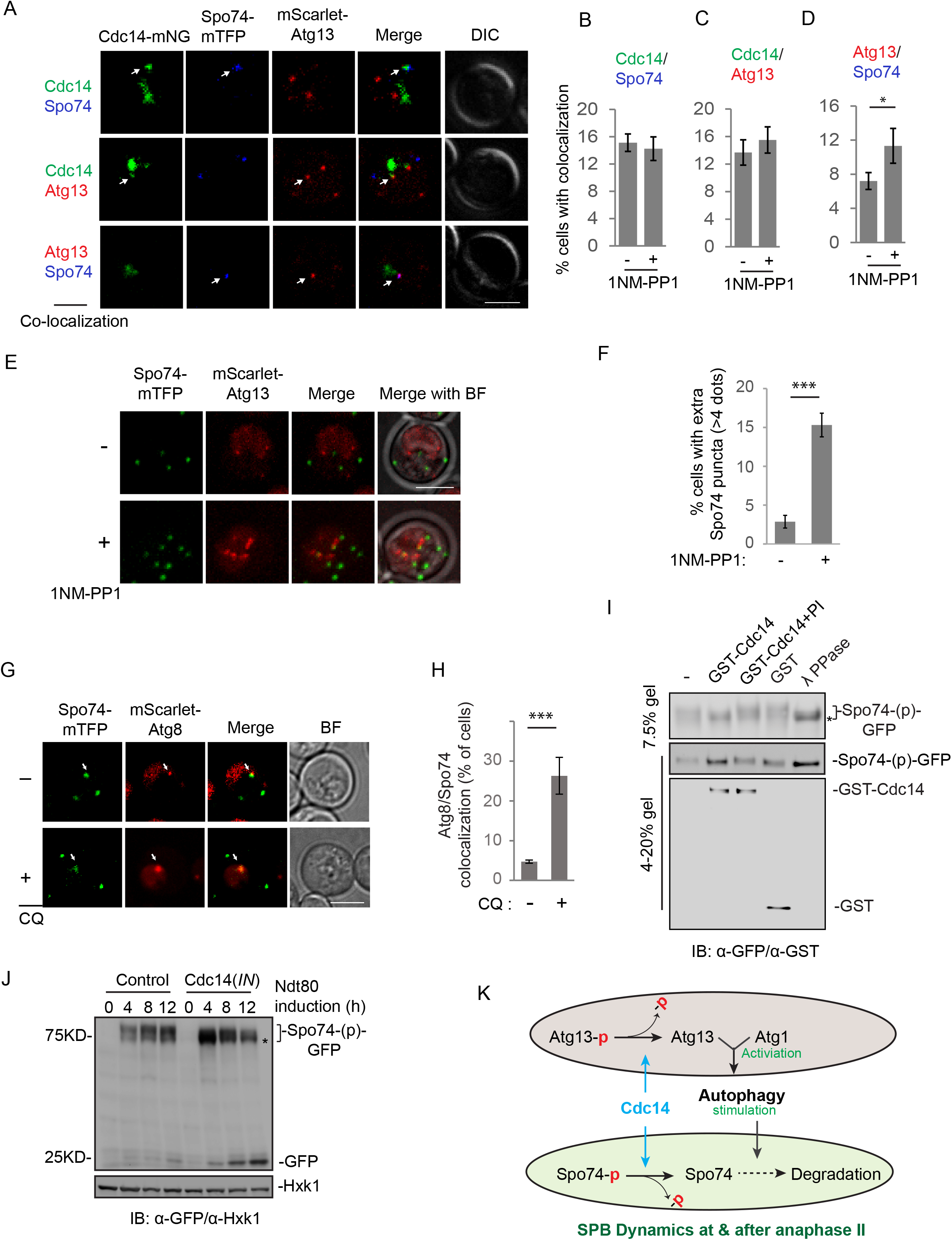
Cdc14 guides autophagy mediated Spo74 degradation at meiosis exit. (A-D) Representative FM images (A) and quantitation (B-D) of colocalization between Cdc14-mNG, Spo74-mTFP, and mScarlet-Atg13 during late meiotic divisions (n≥300 cells, t-test). White arrows indicate colocalization. Scale bar, 5μm. (E & F) Representative FM images (E) and quantification (F) of meiotic cells expressing Spo74-mTFP and mScarlet-Atg13 (n≥300, t-test). Note that extra (> four) Spc74-mTFP puncta formed in the cell with autophagy inhibited by 1NM-PP1. Z-stacked images (5 planes; step size, 0.5µm) were merged in ImageJ. Scale bar, 5µm. (G & H) Representative FM images (G) and quantification (H) of colocalization between Spo74-mTFP and mScarlet-Atg8 during late meiosis & sporulation (6 h after Ndt80 induction, n≥300 cells, t-test). +CQ, 200 µM chloroquine (CQ) treatment for 1 h before imaging. White arrows indicate colocalization. Scale bar, 5μm. (I) 6h after Ndt80 induction, Spo74-GFP in cell lysates were immunoprecipitated (IP), treated as indicated for 20min at 30 °C while immobilized on beads. The samples were then resolved by 7.5% and 4-20% SDS-PAGE followed by IB with indicated antibodies. *, hypo-phosphorylated Spo74-GFP. (J) Whole-cell extracts of synchronized meiotic cells, with or without inducing Cdc14(*IN*), were analyzed by IB with indicated antibodies, showing Spo74-GFP cleavage. Note that Cdc14(*IN*) drives Spo74-GFP band shift to lower position over time and stimulates free GFP generation. *, hypo-phosphorylated Spo74-GFP. (K) Model of Cdc14 stimulating autophagy and Spo74 degradation. At anaphase I & II, Cdc14 dephosphorylates Atg13 to activate atg1 and hence autophagy. Cdc14 further links autophagy with SPB dynamics by dephosphorylating Spo74 to prime its autophagic degradation.

These data suggest that Spo74 is degraded by autophagy during late meiosis. If so, Spo74 should be transiently observed in Atg8-marked autophagosomes. Indeed, we observed colocalization of mScarlet-Atg8 and Spo74-mTFP during meiosis (Fig. 6G & 6H, 4±0.4%). Inhibition of autophagosome–vacuolar fusion by chloroquine (CQ) treatment increased colocalization of mScarlet-Atg8 and Spo74-mTFP by ∼8-fold (Fig. 6G & 6H, 27±4.6%). Consistent with this, IB analysis of late meiosis (and early sporulation) cell lysates revealed that Spo74-GFP degradation resulted in vacuole-resistant GFP, which was repressed upon autophagy inhibition by Atg1-as exposure to 1NM-PP1 (Fig. S6A & S6B). Therefore, we conclude that autophagy degrades Spo74 during late meiosis and sporulation.

CQ treatment led to simultaneous colocalization of Cdc14, Atg8, and Spo74 in 21±2.9% of cells containing colocalized Atg8/Spo74 (Fig. S6C, S6D), suggesting that Cdc14 plays a direct role in autophagy-mediated Spo74 degradation. Intriguingly, inhibition of autophagy by 1NM-PP1 protected hypo-phosphorylated Spo74-GFP to a greater extent than hyper-phosphorylated Spo74-GFP (Fig. S6A), indicating that dephosphorylation primes Spo74 for autophagic degradation. Moreover, colocalization of Cdc14m with Spo74 was dramatically increased relative to Cdc14 (Fig. S6E & S6F, 48%±2%). Furthermore, affinity-purified Spo74-GFP from meiotic cell lysates was efficiently dephosphorylated by recombinant GST-Cdc14 *in vitro* (Fig. 6I), demonstrating that Spo74 is a direct substrate of Cdc14. In addition, Cdc14(*IN*) during meiosis stimulated Spo74 dephosphorylation (Fig. 6J). Together, these data demonstrate that Cdc14 dephosphorylates Spo74 during meiosis. Remarkably, increased Spo74-GFP dephosphorylation by Cdc14(*IN*) correlated with increased autophagic degradation of Spo74-GFP, as indicated by accumulation of free GFP (Fig. 6J), further indicating that dephosphorylated Spo74 by Cdc14 is sensitive to autophagy.

Collectively, these results support a model (Fig. 6K) in which Cdc14 activates Atg1 complex and stimulates the autophagic degradation of meiosis-specific proteins such as Spo74, highlighting how Cdc14 activity licenses the autophagic destruction of developmentally regulated proteins.

## Discussion

In this study, we found that, contrary to previous assumptions (Odle et al., 2020, Wen et al., 2016, Ferder et al., 2019), autophagy is active during meiotic divisions and enables progression into late meiosis and sporulation. In anaphase I and II, Cdc14 stimulates autophagy by dephosphorylating Atg13 to activate Atg1. Cdc14 also promotes autophagic degradation of Spo74, a meiosis-specific SPB component that is essential for the meiosis–sporulation transition. Thus, Cdc14 couples autophagy to the developmentally regulated process of meiotic SPB modulation.

### Autophagy level during meiosis

Cell division, particularly meiosis, is one of the most complex cellular processes, involving dynamic synthesis and destruction of several intracellular structures. Accordingly, cell division requires tightly controlled autophagy. Remarkably, cells with higher levels of autophagic flux during meiotic divisions complete meiosis faster (Fig. 1J, 1K), indicating the importance of Cdc14-mediated autophagy initiation during meiosis. Thus, the Cdc14-dependent stimulation of autophagy initiation during anaphase I and II likely represents a strategy to sequentially boost autophagy above a critical level as cells progress through meiotic divisions. Consistent with this, enhanced autophagy due to expression of Atg13-8SA or Atg13-6SA during meiosis robustly increased sporulation efficiency (Fig. S1F & 5F). This observation, however, raises several interesting questions. For example, what is the range of autophagy levels that benefits meiosis and sporulation the most? Why is autophagy during meiosis not tuned to maximize sporulation, the ultimate purpose of meiosis? One possibility is that high autophagy (and sporulation rate) might reduce the ability of meiotic cells to react to the fluctuating environment in the natural world, e.g., the need to exit meiosis and return to vegetative growth when a nutrient restriction disappears. Thus, a quantitative approach is needed to understand how autophagy levels, with their associated benefits and costs, influence specific cellular processes during meiosis and sporulation.

To fine-tune autophagy levels during meiotic divisions, a cell could use multiple strategies. Little is known about the transcriptional control of meiotic autophagy, but a ribosome profiling study of yeast meiosis reported temporally increased expression of Atg8 (Brar et al., 2012), whose levels influence autophagosome size. Our data demonstrate that Cdc14 controls Atg13 phosphorylation to stimulate autophagy initiation. Given that CDK1 suppresses mitotic autophagy and phosphorylates Atg13 (and Atg1, Atg14 etc), Cdc28 (CDK1) is likely to counteract Cdc14 in the regulation of autophagy during yeast meiosis. Although the coordination between Cdc28 and Cdc14 in autophagy control is yet to be established, this phosphorylation-based mechanism of regulating meiotic autophagy might include factors other than Cdc28 and Cdc14, e.g., the meiosis-specific kinase Ime2 or the TORC1 kinase and the PP2 family protein phosphatases. Although TORC1 inhibition is a prerequisite for yeast meiosis entry, the low remaining levels of TORC1 activity during meiosis might be sufficient to regulate Atg13. In support of this notion, we found that rapamycin upregulated autophagy initiation during meiosis, as indicated by increases in Atg1 (Fig. S2D) and Atg13 puncta formation (Fig. S2E). In addition to Cdc14, PP2A and PP2C stimulate autophagy under stress conditions, e.g., starvation and genotoxin treatment (Yeasmin et al., 2016, Memisoglu et al., 2019), respectively. Thus, the potential contributions of kinases (e.g., Ime2, Cdc28, and TORC1) and phosphatases (e.g., PP2A, and PP2C) in modulating autophagy during meiosis need to be further investigated. We speculate that bursts in phosphorylation/dephosphorylation orchestrated by these conserved kinases and phosphatases during meiosis integrate inactivation and activation of autophagy during the progression of meiosis and sporulation.

### Cdc14 as a regulator of autophagy

Cdc14 is a dual phosphatase with a structurally defined binding preference for a PxL motif (Kataria et al., 2018). Our data demonstrate that the PxL3 motif of Atg13 is required for Cdc14 docking on the PAS, suggesting that Atg13 is the main binding partner of Cdc14 at the PAS. Intriguingly, the PxL3 motif in the flanking IDR of yeast Atg13, which has access to approximately five SPs (Cdc14 dephosphorylation sites), is conserved across species, suggesting that the Cdc14–Atg13– autophagy axis revealed in this study might be conserved in higher eukaryotes.

Atg13 might not be the only target of Cdc14 involved in the regulation of autophagy. The PAS is a structurally undefined meshwork of the Atg1 protein complex, PI3K complex, Atg2–Atg18 complex, and Atg9 vesicles. In addition to Atg13, several PAS-localized Atg proteins (e.g., Atg1 and Atg18) also have PxL sites in their LCD regions that are accessible to Cdc14 docking and flanked by several SP sites. Hence, it is possible that docking of the Cdc14 homodimer on the PAS is a consequence of the cooperation of multiple PAS components and that the resultant Cdc14-mediated dephosphorylation might go beyond Atg13. Moreover, the fact that the mid-late autophagosome marker Atg8 colocalizes with Cdc14 indicates that Cdc14 might interact and function in mid-late stages of autophagosome biogenesis, likely independent of Atg13 binding or dephosphorylation. (Note that Atg8 colocalizes with Cdc14 to a lesser extent than Atg13 or Atg1, suggesting that Cdc14 remains associated with the growing/mature autophagosome.) Further studies are required to reveal whether and in what manner Atg1 and Atg18, or other Atg proteins, are targets of Cdc14 phosphatase activity.

Under our experimental conditions, autophagy triggered in interphase cells by starvation did not stimulate Cdc14 cytosolic release or Cdc14/Atg13 colocalization (Fig. S3G). Moreover, we did not observe Cdc14–Atg13 colocalization during mitotic anaphase (Fig. S3G). Thus, Cdc14 is likely a meiosis-specific autophagy regulator. It remains unknown whether meiotic factor(s) assists Cdc14 in stimulating autophagy during anaphase I and II, and whether mitotic mechanisms prevent Cdc14 from engaging in autophagy initiation during mitotic anaphase.

### Autophagy substrates during meiotic cell division

It is not clear whether the restriction of Cdc14’s effect on autophagy to meiotic anaphase reflects that autophagy is tailored for each meiotic stage and specifically needed during anaphases. We previously identified Rim4 (Wang et al., 2020b) as a meiosis-specific substrate of autophagy at anaphase I; in this study, we identified Spo74 as a meiosis-specific substrate of autophagy at anaphase II. Rim4 assembles into aggregates at early meiotic stages, sequestering a subset of mid-late meiotic transcripts to prevent their translation. Therefore, timely translation of Rim4-sequestered transcripts relies on rapid removal of Rim4 aggregates at the onset of meiosis II, which is accomplished by autophagy rather than proteasomal degradation. On the other hand, Spo74 in SPB is required for prospore membrane biogenesis by facilitating docking and fusion of Golgi-derived vesicles (Neiman, 2011, Nakanishi et al., 2006, Nickas et al., 2003). The biochemistry of the SPB is akin to some protein aggregates, in that it involves the oligomerization or self-assembly of several protein components onto a lattice; in the case of the S. *cerevisiae* SPB, the lattice is provided by Spc42 oligomerization (Jaspersen, 2021), with Spo74 residing on the cytosolic side of the giant >0.5 GDa complex. Extra Spo74-containing SBPs form in the absence of autophagy, indicated by extra dots of Spo74 (Fig. 6E & 6F), Mpc54, Spo21, and Spc42 in the cell at the end of meiosis (Wang et al., 2020b). These supernumerary SBPs cause abnormal chromosome segregation and aberrant prospore formation (Wang et al., 2020b). These observations highlight the importance of tight control of SPB numbers and suggest that this control is likely to be achieved by autophagy-mediated degradation of SPB proteins, including Spo74. It is worth noting that loss of autophagy does not affect SPB numbers in mitosis. Thus, the Cdc14-promoted upregulation of autophagy at anaphase I and II suggests the existence of meiotic autophagy substrates that must be identified before we can fully understand how autophagy affects meiotic translation via Rim4, and how meiotic SPB number is guided by autophagy.

Why does meiosis use Cdc14 to facilitate autophagy-mediated Spo74 degradation? A key function of kinase signaling in meiotic divisions is the establishment of a tightly scheduled sequence of interdependent cellular events. Intriguingly, Spo74 dephosphorylated by Cdc14 is more sensitive to autophagy (Fig.6J & S6A). It is well established that phosphorylation/dephosphorylation of SPB components affects SPB assembly and disassembly (Jaspersen et al., 2004). Cdc14 colocalizes with meiotic SPBs during the anaphases. Thus, dephosphorylation of Spo74 by Cdc14 could be spatially constrained to the SPBs. Such a localized mechanism might allow the establishment of dependent relationships between Spo74 dephosphorylation and its autophagic degradation. If so, SPB-localized Cdc14 could guide the autophagic machinery to targets on the SPB, such as Spo74. One example of such spatially constrained dephosphorylation involves PP2A, which localizes its activity to the region of centromeres to retain meiotic cohesion during and after meiosis I (Riedel et al., 2006). How autophagy regulates SPB dynamics by discriminating the phosphorylation state of specific SPB component(s), including Spo74, remains an important topic for future research.

## Supporting information

Movie S1 A representative cell undergoing meiosis

## Acknowledgments

We thank Beth Levine, the UTSW Center for Autophagy Research, the UTSW Live Cell imaging facility, and Benjamin Tu for imaging and experimental support. We thank Vladimir Denic, Soni Lacefield for reagents and protocols. We thank B. Levine, J. Seemann, J. Friedman, S. Lacefield, M. Henne, and members of the Wang lab for comments on the manuscript. This work was supported by a grant from the NIH to F.W. (R01GM133899) and from the Welch Foundation to F.W. (I-2019-20190330), as well as funding from Nancy Cain and Jeffrey A. Marcus Scholar in Medical Research, in Honor of Dr. Bill S. Vowell, to F.W. and the National Institute of General Medical Sciences of the National Institutes of Health (K99GM13548) to O.A.M.

## Author contributions

Conceptualization, F.W. and W.F.; Methodology, F.W., W.F., and O.A.M.; Investigation, F.W., W.F., O.A.M., and S.Q.; Writing – Original Draft, F.W.; Writing – Review & Editing, F.W., O.A.M., and W.F.; Funding Acquisition, F.W.; Supervision, F.W.

## Supplemental Figures

**Figure S1.**
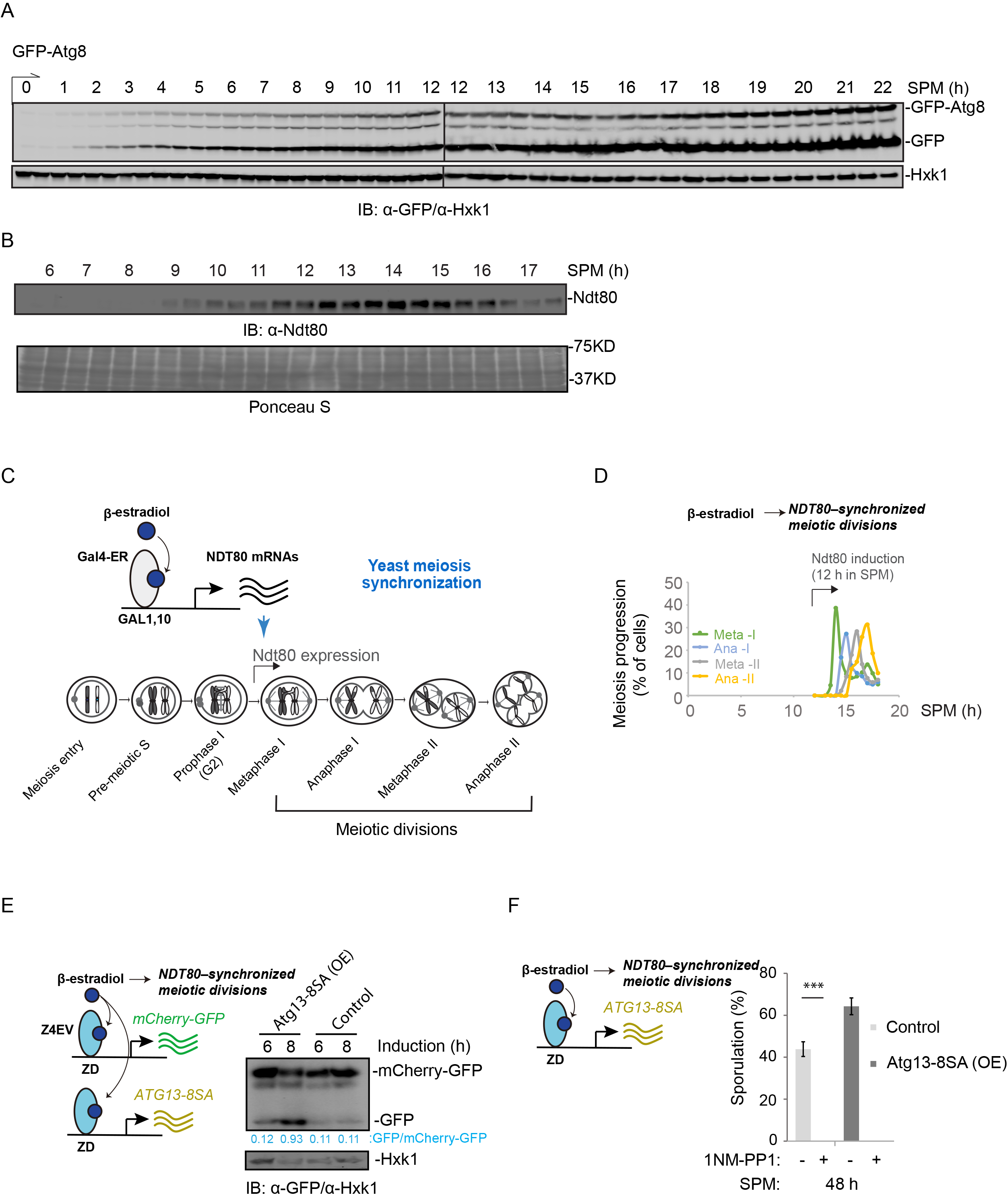
Autophagy levels influence meiosis and sporulation. (A) Whole-cell extracts derived from cells during meiosis were treated as diagrammed in (1A) and analyzed by immunoblotting (IB) with indicated antibodies. GFP-Atg8 expression was induced upon SPM incubation (0 h). (B) Top, IB (α-Ndt80) showing Ndt80 expression in the same samples prepared as in (S1A). Bottom, proteins transferred to the blotting membrane were stained with Ponceau S before IB. (C) Schematic of Gal1-Ndt80–synchronized meiotic divisions. The *GAL1,10* promoter replaces the *NDT80* promoter at the endogenous locus to enable β-estradiol-inducible expression by the chimeric transcription factor Gal4-ER. Unless indicated otherwise, 1μM β-estradiol was added to meiotic cells after 12 h in SPM to synchronize the entry of meiotic divisions, depicted as t=0 h of Ndt80 induction. (D) Graph of Gal-Ndt80–synchronized meiotic progression showing the percent of cells at metaphase I, anaphase I, metaphase II, and anaphase II at indicated time points. Cells derived from indicated time points were fixed and proceeded with immunofluorescence staining (IF: α-tubulin Alex 546) as described in the method. (E) IB of meiotic cell lysates with indicated antibodies, showing the effect of Atg13-8SA (OE) on mCherry-GFP cleavage at indicated time after Ndt80 induction. Left, schematic of simultaneously induced expression of Atg13-8SA and mCherry-GFP during meiotic divisions by 1µM β-estradiol, which also induced Ndt80 expression to synchronize the entry of meiotic divisions. (F) Graph of sporulation efficiency, showing the effect of Atg13-8SA (OE) induction and 1NM-PP1 treatment during meiotic divisions on sporulation. Percent of cells with spores after 48 h in SPM are shown. (n≥300 cells from three replicate experiments; t-test). Left, schematic of induced expression of Atg13-8SA and Ndt80 by 1µM β-estradiol. 5uM 1NM-PP1, if applied, was applied at the same time.

**Figure S2.**
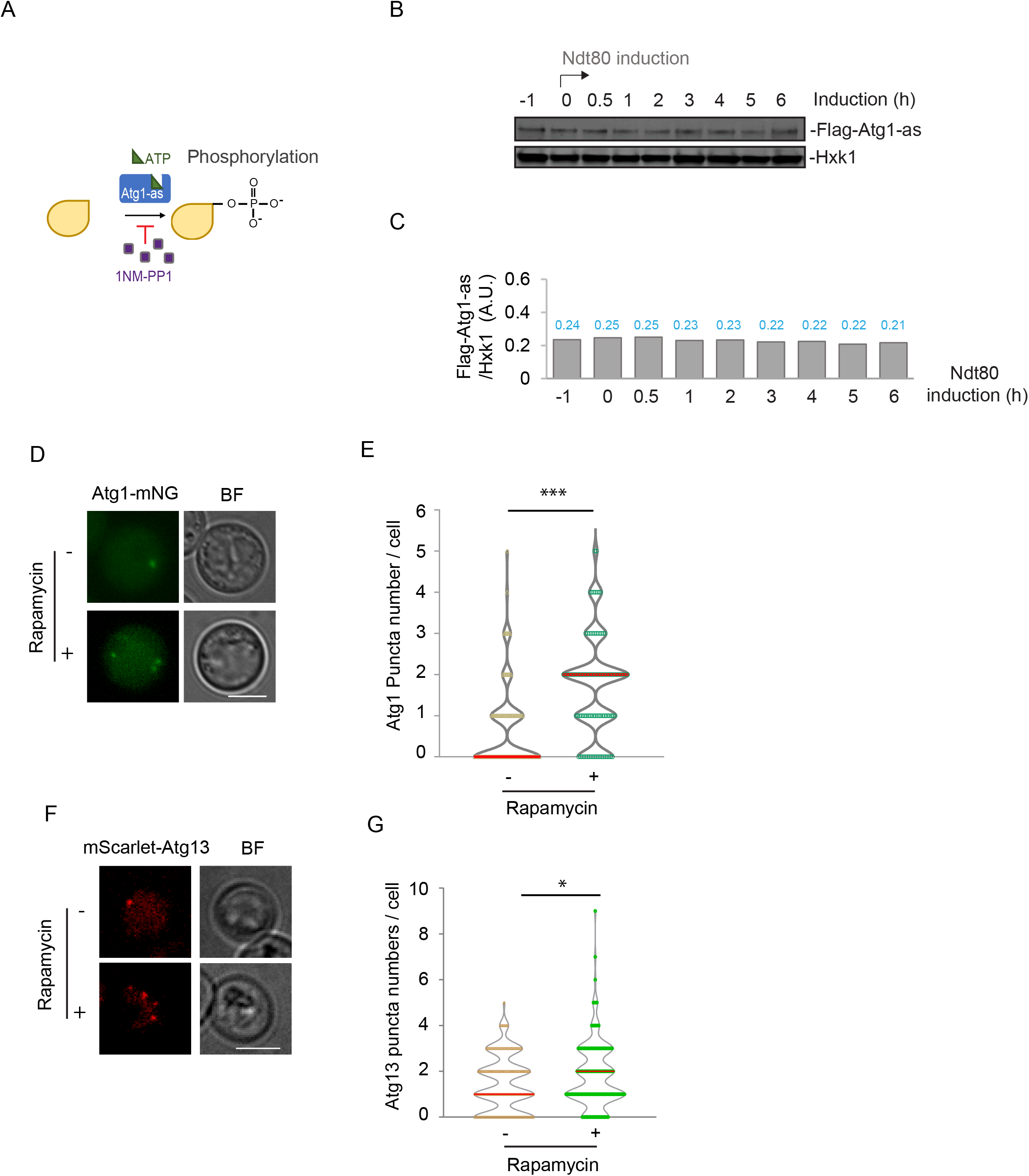
Autophagy initiation during meiotic division can be stimulated by TORC1 inactivation. (A) Schematic of the chemical-genetic strategy for inhibiting the kinase activity of Atg1-as. Atg1 was genetically mutated to its *Shokat* allele, Atg1-M102G, with the bulky ‘‘gatekeeper residue’’ in the ATP-binding pocket of Atg1 replaced to create a functional ATP-analog-sensitive allele of Atg1 (Atg1-as). 1NM-PP1, a membrane-permeable ATP analog can specifically inhibit Atg1-as kinase activity in a live cell. (B & C) Whole-cell extracts derived from meiotic cells were analyzed by immunoblotting (IB) with α-FLAG and α-Hxk1 antibodies. Graph (C) shows the quantification of (B). Numbers on top (blue) show values of Atg1 protein IB intensity in arbitrary units, normalized by Hxk1 IB intensity. (D & E) Representative FM images (D) and quantification (E) of Atg1-mNG puncta numbers per cell with or without Rapamycin treatment during synchronized meiotic divisions (n=100 cells, t-test). Red line, median. FM images were captured and quantified after 3 h of Rapamycin (0.2 µM) treatment and Ndt80 induction. (F & G) Representative FM images (F) and quantification (G) of mScarlet-Atg13 puncta numbers per cell with or without Rapamycin treatment during synchronized meiotic divisions (n=150 cells, t-test). Red line, median. Experimental condition was as described (D & E).

**Figure S3.**
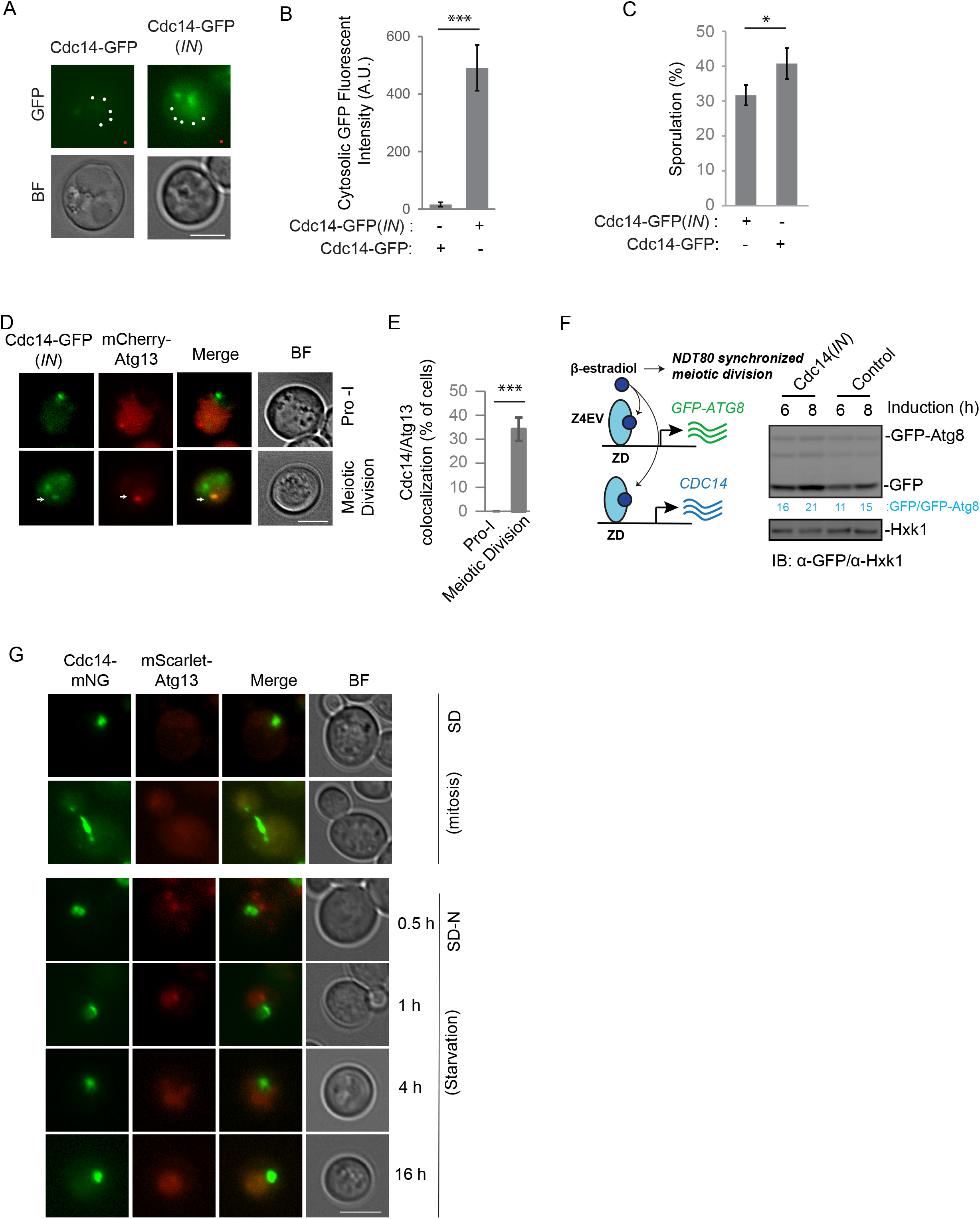
Cdc14 stimulates autophagy during meiosis. (A & B) FM analysis of Cdc14-GFP and Cdc14-GFP (*IN*) at Anaphase II (A) and the quantitation of GFP fluorescence signal (B). The fluorescence intensity measured at the white dots was normalized to the background signal (red dot) and shown as an average arbitrary unit (A. U.) (n=10 cells, 5 white dots/cell, t-test). Scale bar, 5µm. (C) Percent of cells showing sporulation after 48 h in SPM with indicated conditions (n≥300 cells; t-test). Cd14-GFP(*IN*) was induced by 1µM β-estradiol at t=0 h, if needed. (D & E) FM analysis of mCherry-Atg13 colocalization with Cdc14 GFP (*IN*) during prophase I and meiosis divisions. Shown are representative FM images (D) and quantitation (E) of mCherry-Atg13/Cdc14 GFP (*IN*) colocalization (n≥300 cells; t-test). Scale bar, 5µm. (F) IB analysis with indicated antibodies, showing the change of GFP-Atg8 processing in response to Cdc14(*IN*). Left, schematic of simultaneously induced expression of Cdc14, GFP-Atg8, and Ndt80. (G) Representative FM images showing Cdc14-mNG and mScarlet-Atg13 fluorescence signal in cells under vegetative growth condition (SD medium, log-phase cells) or nitrogen starvation condition (SD-N medium). Scale bar, 5µM. n≥300 cells.

**Figure S4.**
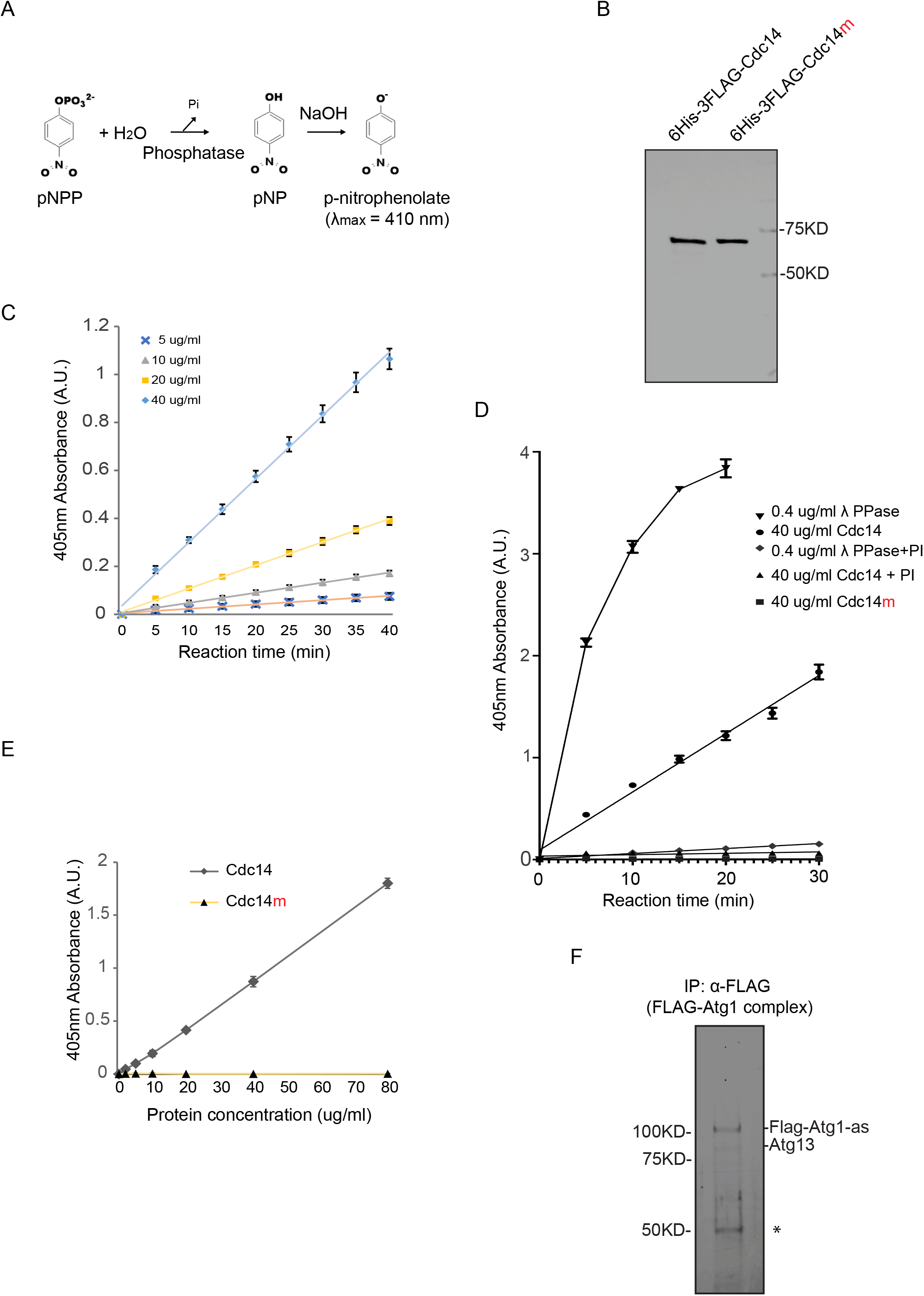
Recombinant Cdc14 carries phosphatase activity *in vitro*. (A) Schematic of the colorimetric phosphatase activity assay measured spectrophotometrically at 410 nm by determining the amount of pNP produced following the hydrolysis of Pi from pNPP. The addition of NaOH after a specified assay time (e.g., 10 min) serves to stop the phosphatase reaction while simultaneously converting the product p-nitrophenol into the yellow-colored p-nitrophenolate (λmax = 410 nm). (B) The indicated proteins were prepared from *E. Coli*, as detailed in Experimental Procedures for recombinant protein expression and purification and analyzed by SDS-PAGE followed by Coomassie Blue staining. (C) Phosphatase activity of recombinant 6His-3FLAG-Cdc14 protein with indicated concentrations was assayed as in part (S4A). Shown are the average and standard deviation, from three replication experiments, of p-nitrophenolate absorbance at 405 nm plot to incubation time. (D) Phosphatase activity of 40 µg/ml recombinant 6His-3FLAG-Cdc14, 40 µg/ml 6His-3FLAG-Cdc14m or 0.4 µg/ml λ phosphatase was assayed in the presence or absence of phosphatase inhibitor (PI) as in part (S4A), and shown as (S4C). Note that PI almost abolished phosphatase activity in 6His-3FLAG-Cdc14 and λ phosphatase. (E) Phosphatase activity of recombinant 6His-3FLAG-Cdc14 and 6His-3FLAG-Cdc14m protein was assayed for 30 min as in part (S4A). Shown are the average and standard deviation, from three replication experiments, of p-nitrophenolate absorbance at 405 nm plot to protein concentration. Note that 6His-3FLAG-Cdc14m yielded no detectable absorbance over the background. (F) Immunoprecipitation (IP: α-FLAG) of Atg1 (FLAG-Atg1-as) complex from cell lysate derived from the *FLAG-Atg1-as* cells arrested at Prophase I. Following washing, the Atg1 (FLAG-Atg1-as) complex was eluted from the resin with FLAG peptide. Eluted proteins were resolved by SDS-PAGE and visualized by Sypro Ruby staining. Flag-Atg1-as protein identity was confirmed by IB. Atg13 was putatively assigned based on its size. *, Igg antibody.

**Figure S5.**
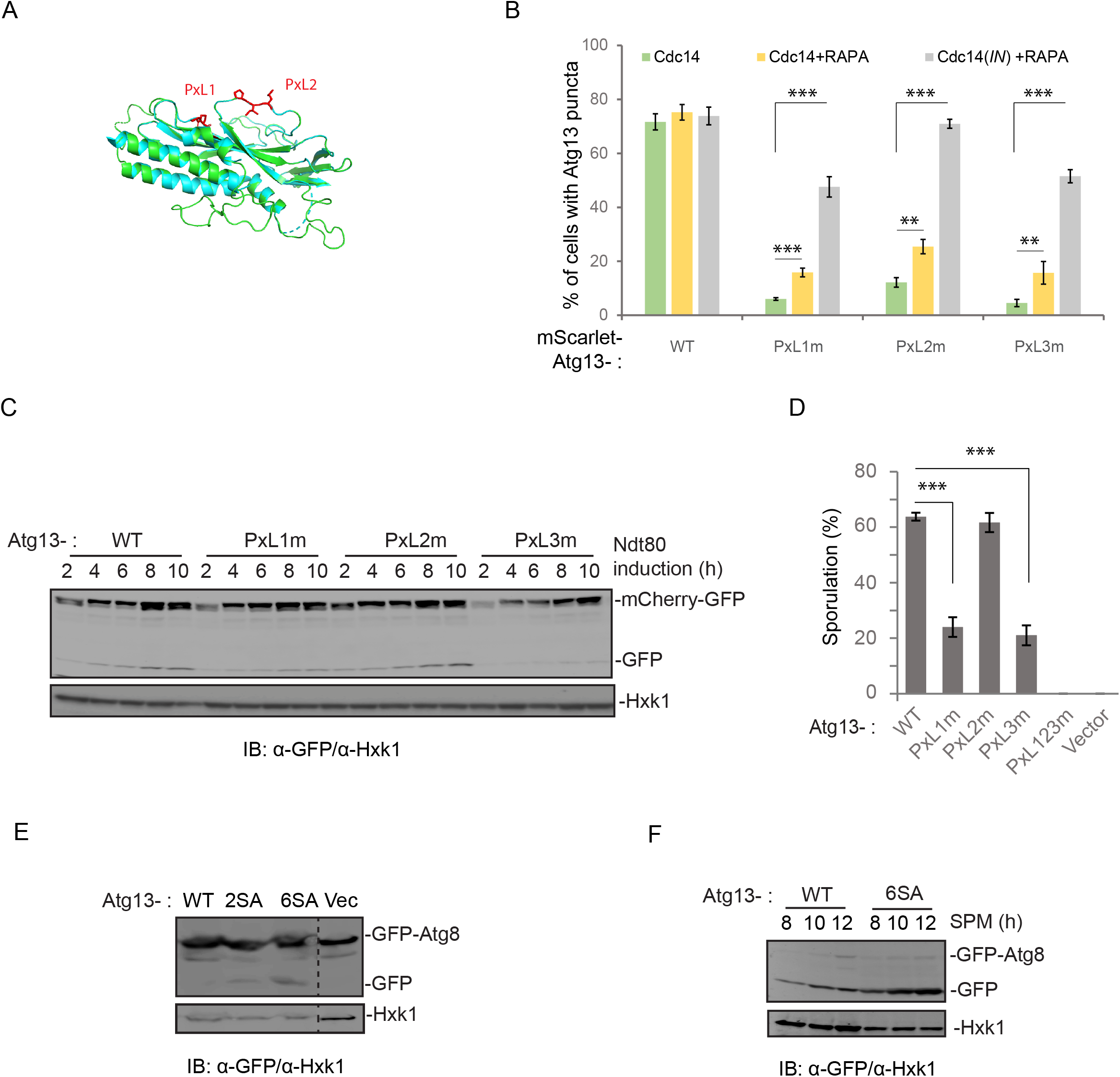
Two PxL motifs in Atg13 are critical for the function of Atg13 in autophagy. (A) A PyMOL cartoon representation of the homology model of budding yeast (S. *cerevisiae*) Atg13 residues 7 to 267 (Green) made by SWISS-MODEL homology modeling (Waterhouse et al., 2018) superimposed on the cartoon of the crystal structure of *Lachancea thermotolerans* Atg13 HORMA domain (blue) (PDB: 4J2G). PxL1 and PxL2 sites mutated in this study are indicated as sticks in red. (B) Graph showing the percent of cells with puncta of Atg13 (mScarlet-Atg13) or its variants, with indicated conditions, during the meiotic divisions. Cells with mScarlet-Atg13 puncta were quantified by FM analysis at t=6 h after Ndt80 induction (n≥300, t-test). Rapamycin, if applied, was added to growth medium 2 h (t=-2 h) before Ndt80 induction. (C) IB of cell extracts from meiotic cells with indicated antibodies. The mCherry-GFP (p*ZD: mCherry-GFP*) and Ndt80 were simultaneously induced by 1µM β-estradiol. (D) Atg13 variants carried by pRS303 or empty vector were introduced into *atg13Δ* cells, in which sporulation was triggered. Shown is the percent of cells showing sporulation in 48 h. (n ≥300, ***, p≤0.001, t-test). (E) Atg13 variants carried by pRS303 or empty vector were introduced into *atg13Δ* cells, in which GFP-Atg8 (pZD: GFP-Atg8) was induced by β-estradiol for 4 h during vegetative growth (log-phase). The whole-cell lysates were resolved by SDS-PAGE and analyzed by IB with indicated antibodies. 2SA, Atg13-S129A-S454A; 6SA, Atg13-S129A-S454A-S535A-S541A-S646A. Note that the GFP-Atg8 processing levels in 2SA and 6SA cells increased, indicated by increased free GFP generation normalized by GFP-Atg8. (F) Atg13 or Atg13-6SA carried by pRS303 was introduced into *atg13Δ* cells, which harbor GFP-Atg8 (pZD: GFP-Atg8). After initiating sporulation for 6 h, GFP-Atg8 expression was induced in these strains, and cells were harvested at indicated time points. The whole-cell lysates were resolved by SDS-PAGE and analyzed by IB with indicated antibodies. Note that free GFP generation increased in Atg13-6SA cells.

**Figure S6.**
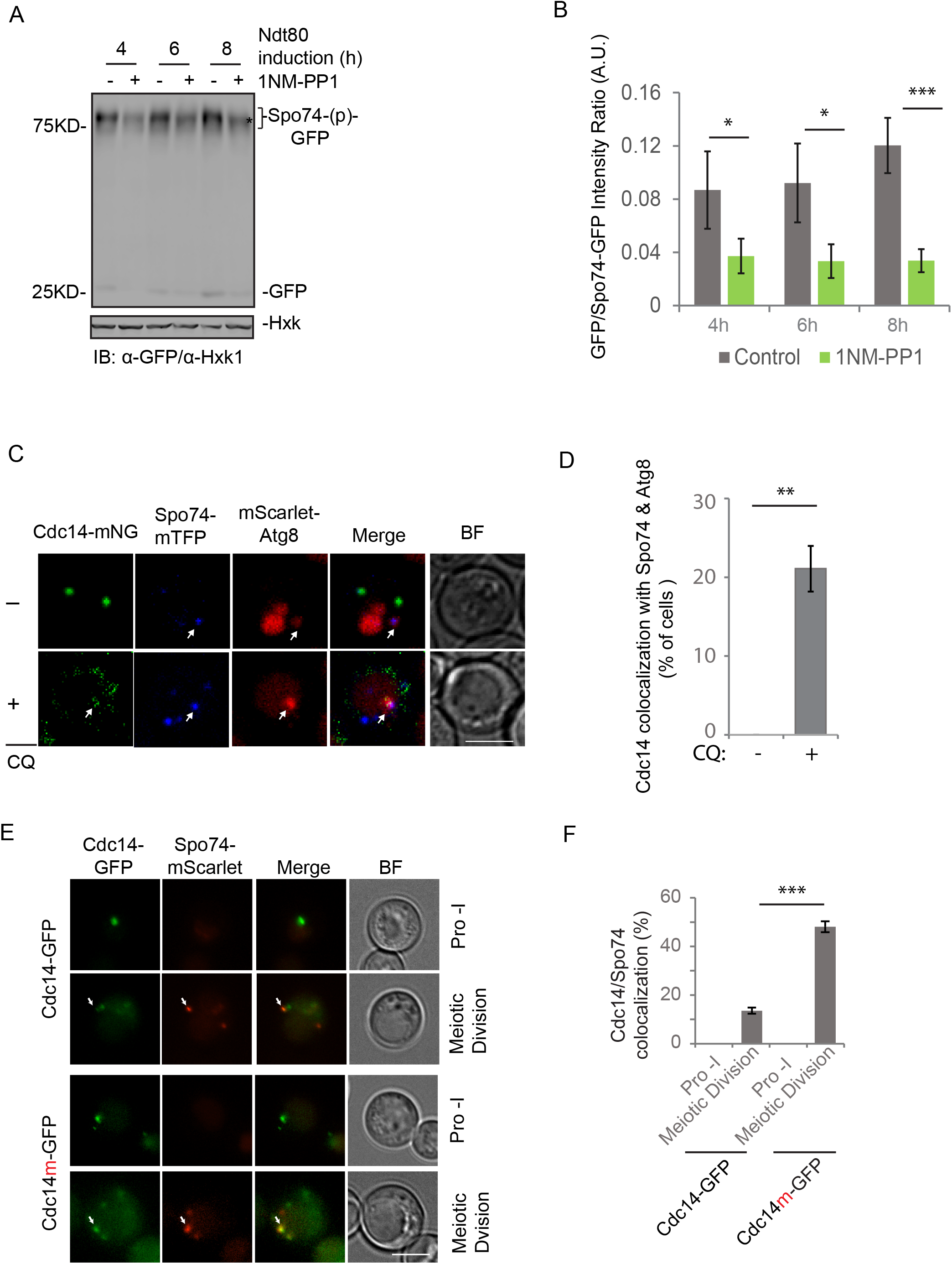
Cdc14 guides autophagy mediated Spo74 degradation at meiosis exit. (A) IB of synchronized meiotic cell lysates with indicated antibodies, showing Spo74-GFP cleavage. 1NM-PP1 was applied when Ndt80 was induced by β-estradiol (t=0). Note that 1NM-PP1 caused accumulation of hypo-phosphorylated Spo74-GFP. *, hypo-phosphorylated Spo74-GFP. (B) Quantification of (A) showing average GFP/Spo74-GFP ratio with standard deviation. IB images from 3 independent experiments were quantified by Bio-Rad Image Lab software. (*, p≤0.01, ***, p≤0.001, t-test) (C & D) Representative FM images (E) and quantification (F) of Cdc14-mNG simultaneous colocalization with Spo74-mTFP & mScarlet-Atg8 during late meiosis (6 h after Ndt80 induction). In total, 100 cells showing Spo74-mTFP/mScarlet-Atg8 colocalization were quantified (Mann-Whitney U-test). +CQ, 200 µM CQ treatment for 1 h before imaging. Scale bar, 5μm. (E & F) Representative FM images (C) and quantification (D) of cells during Prophase-I and meiotic divisions, showing colocalization of Cdc14-GFP/Cdc14m-GFP and Spo74-mScarlet. Cell expresses Cdc14-GFP or Cdc14m-GFP under their endogenous promoters from pRS304 vector, along with genomic tagged Spo74-mScarlet. Note that *cdc14* genes at their genome locus are intact. (C) White arrows indicate colocalization. Scale bar, 5µm. (D) Only cells that carry Spo74-mScarlet puncta were analyzed (n≥300; ***, p≤0.001, t-test)

**Table S1.**
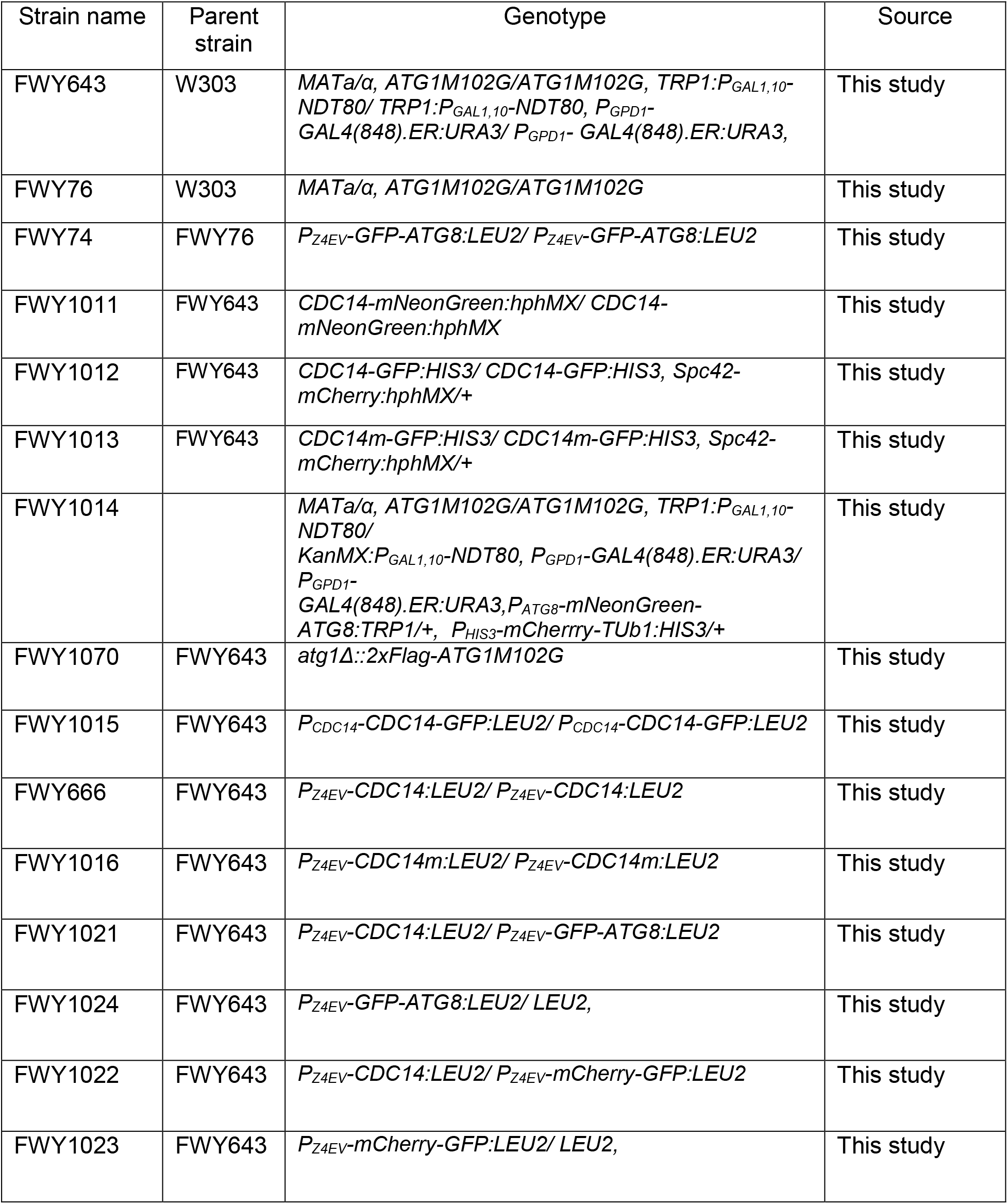
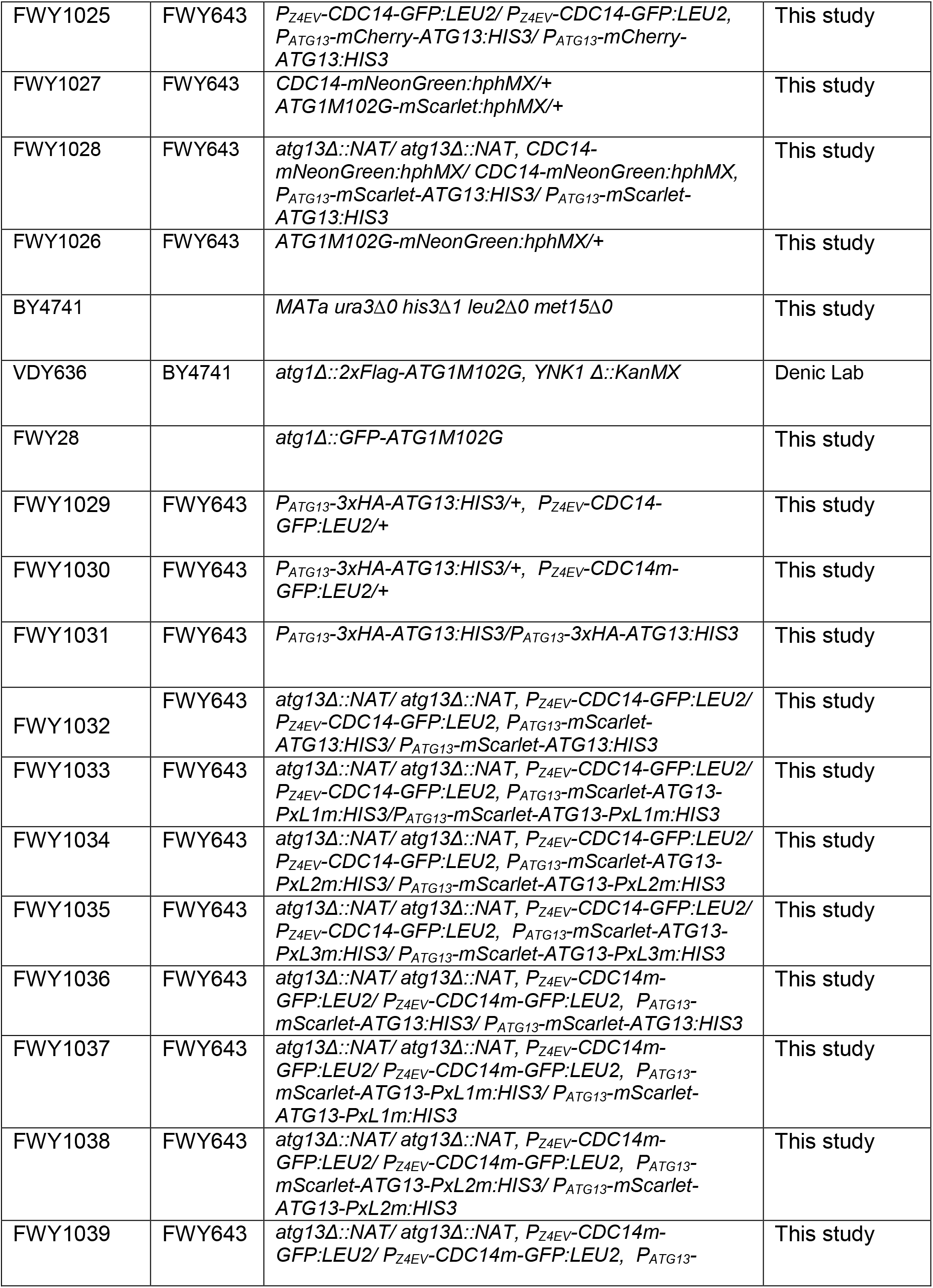
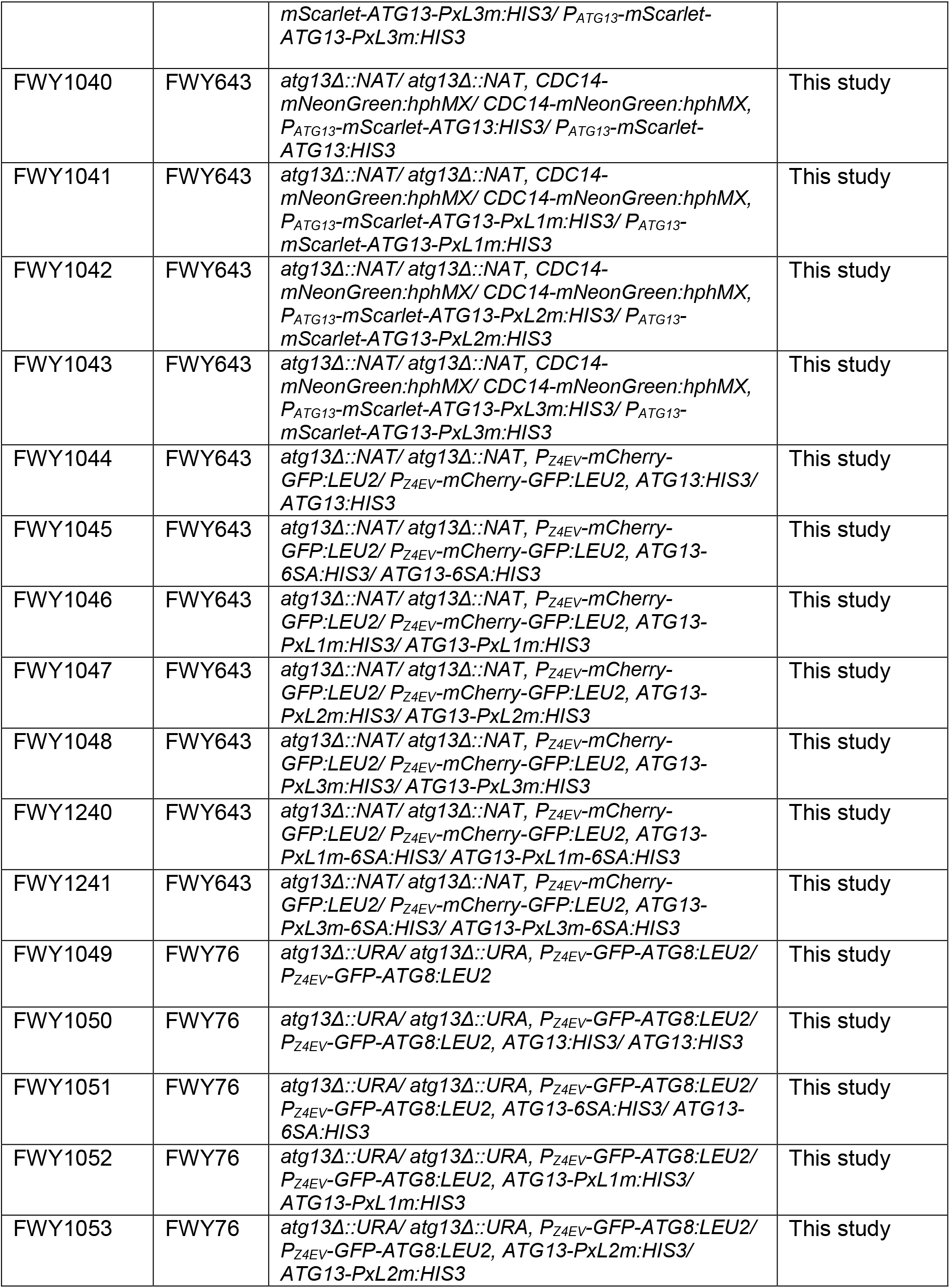
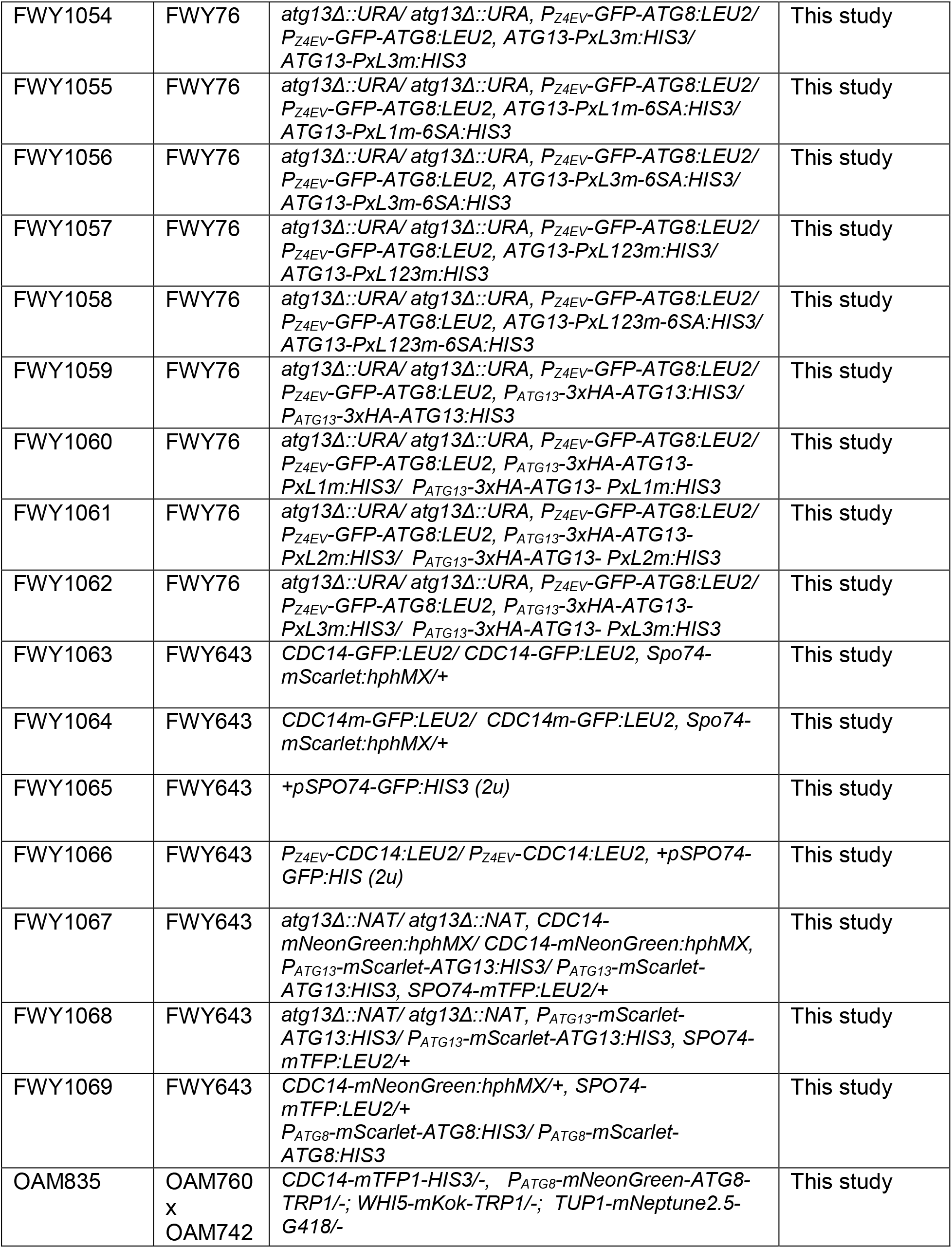
Strains used in this study. All strains are derivatives of W303 (*ade2-1 his3-11,15 leu2-3,112 trp1-1 ura3-1 can1-100*)

**Table S2.** Plasmids used in this study.

| Plasimd | Vector | Mutations | Source |
| --- | --- | --- | --- |
| *P<sub>GPD1</sub>-GAL4(848).ER:URA3* | pRS306 |  | Wang *et al*, 2020 |
| *P<sub>Z4EV</sub>-GFP-ATG8:LEU2* | pNH605 |  | Wang *et al*, 2020 |
| *P<sub>ATG8</sub>-2xGFP-ATG8:HIS3* | pRS303 |  | This study |
| *P<sub>ATG8</sub>-mNeonGreen-ATG8:LEU2* | pRS305 |  | This study |
| *P<sub>ATG8</sub>-mNeonGreen-ATG8:TRP1* | pRS304 |  | Arguello-Miranda *et al.*, 2021 |
| *P<sub>Z4EV</sub>-mCherry-GFP:LEU2* | pNH605 |  | This study |
| *P<sub>Z4EV</sub>-CDC14:LEU2* | pNH605 |  | This study |
| *P<sub>Z4EV</sub>-CDC14m:LEU2* | pNH605 | C283S | This study |
| *P<sub>Z4EV</sub>-CDC14-GFP:LEU2* | pNH605 |  | This study |
| *P<sub>Z4EV</sub>-CDC14m-GFP:LEU2* | pNH605 |  | This study |
| *P<sub>CDC14</sub>-CDC14-GFP:LEU2* | pRS305 |  | This study |
| *P<sub>CDC14</sub>-CDC14m-GFP:LEU2* | pRS305 | C283S | This study |
| *P<sub>T7</sub>-6xHIS-3xFLAG-CDC14* | pET-28a |  | This study |
| *P<sub>T7</sub>-6xHIS-3xFLAG-CDC14m* | pET-28a | C283S | This study |
| *P<sub>T7</sub>-GST-CDC14* | pGEX-6P-1 |  | This study |
| *P<sub>ATG13</sub>-ATG13:HIS3* | pRS303 |  | This study |
| *P<sub>ATG13</sub>-ATG13-2SA:HIS3* | pRS303 | S129A, S454A | This study |
| *P<sub>ATG13</sub>-ATG13-6SA:HIS3* | pRS303 | S129A, S348A, S454A, S535A, S541A, S646A | This study |
| *P<sub>ATG13</sub>-ATG13-PxL1m:HIS3* | pRS303 | P76A, L78G | This study |
| *P<sub>ATG13</sub>-ATG13-PxL2m:HIS3* | pRS303 | P207A,L209G | This study |
| *P<sub>ATG13</sub>-ATG13-PxL3m:HIS3* | pRS303 | P476A,L478G | This study |
| *P<sub>ATG13</sub>-ATG13-PxL1m-6SA:HIS3* | pRS303 | P76A, L78G, S129A, S348A, S454A, S535A, S541A, S646A | This study |
| *P<sub>ATG13</sub>-ATG13-PxL3m-6SA:HIS3* | pRS303 | P476A,L478G, S129A, S348A, S454A, S535A, S541A, S646A | This study |
| *P<sub>ATG13</sub>-3xHA-ATG13:HIS3* | pRS303 |  | This study |
| *P<sub>ATG13</sub>-3xHA-ATG13-PxL1m:HIS3* | pRS303 | P76A,L78G | This study |
| *P<sub>ATG13</sub>-3xHA-ATG13-PxL2m:HIS3* | pRS303 | P207A,L209G | This study |
| *P<sub>ATG13</sub>-3xHA-ATG13-PxL3m:HIS3* | pRS303 | P476A,L478G | This study |
| *P<sub>ATG13</sub>-mScarlet-ATG13:HIS3* | pRS303 |  | This study |
| *P<sub>ATG13</sub>-mScarlet-ATG13-PxL1m:HIS3* | pRS303 | P76A,L78G | This study |
| *P<sub>ATG13</sub>-mScarlet-ATG13-PxL2m:HIS3* | pRS303 | P207A,L209G | This study |
| *P<sub>ATG13</sub>-mScarlet-ATG13-PxL3m:HIS3* | pRS303 | P476A,L478G | This study |
| *P<sub>SPO74</sub>-SPO74-GFP:LEU2:2μm* | pRS425 |  | Wang *et al*, 2020 |
| *P<sub>SPO74</sub>-SPO74-mTFP:LEU2* | pRS305 |  | This study |
| *P<sub>ATG8</sub>-mScarlet-ATG8:HIS3* | pRS303 |  | This study |

**Movie S1:** Autophagy and meiotic progression at the single cell level. A representative cell undergoing meiosis expressing the Cdk1-activity marker Whi5-mKOκ, the autophagy reporter mNG-Atg8, the phosphatase Cdc14-mTFP1, and nuclear marker Tup1-mNeptune2.5. Experimental sampling rate was 6 min, 3 frames/sec.

## Methods

### Saccharomyces cerevisiae Strain Construction

*Saccharomyces cerevisiae* strains are derivatives of W303 (*ade2-1 his3-11,15 leu2-3,112 trp1-1 ura3-1 can1-100*) (Table S1). Unless otherwise indicated, the analog sensitive version of Atg1(Atg1-as) was created by gatekeeper residue change (M102G) as described earlier (Blethrow et al., 2004, Kamber et al., 2015), and introduced into the background of listed strains to allow conditional autophagy inhibition by 1NM-PP1.

Deletion strains were constructed in a parent background by PCR-mediated knock-out with one of the following drug resistance or prototrophic marker cassettes: pFA6a-kanMX6/pFA6a-NAT (Longtine et al., 1998), pKlURA/pCgHIS (Goldstein and McCusker, 1999). C-or N-terminal tagging at the endogenous genomic locus was introduced by PCR-mediated epitope tagging as previously described (Denic and Weissman, 2007). Specifically, strains with *GAL-NDT80 GAL4-ER* were constructed by replacing the endogenous *NDT80* promoter with the inducible *GAL1,10* promoter as described (Carlile and Amon, 2008). Strains with β-estradiol-inducible *GFP-ATG8*, *Cdc14-GFP*, *mCherry-GFP*, or *Atg13 8SA* transgene were made by homologous recombination of a *Z4EV* expression cassette with a Z4EV-driven (*ZD*) promoter (McIsaac et al., 2013) upstream of the *GFP-ATG8*, *Cdc14-GFP mCherry-GFP*, or *Atg13 8SA* open reading frame at the *LEU2* locus. Tub1-mCherry, Cdc14-mTFP, Cdc14-mNeonGreen, Atg1-as-mNeonGreen, Atg1-as-mScarlet, GFP-Atg1-as, Flag-Atg1-as, and Spo74-mScarlet were generated at their endogenous genome locus by transformation with PCR tagging cassettes of indicated C-terminal tags using the PEG/lithium acetate yeast transformation kit (Millipore-Sigma, YEAST1). Plasmids with 2XGFP-Atg8 in pRS303, mNeonGreen-Atg8 in pRS305, Cdc14-GFP in pRS305, Cdc14m-GFP in pRS305, mCherry-Atg13 in pRS303, 3HA-Atg13 in pRS303, mScarlet-Atg13 in pRS303, mScarlet-Atg8 in pRS303 or Spo74-mTFP in pRS303, driven by their endogenous promoters, were integrated into the genomic locus of *HIS3/LEU2* (Wang et al., 2020).

The OAM835 strain was constructed for microfluidics experiments, which resulted from mating haploids obtained by tetrad dissection or by transformation with PCR tagging cassettes or plasmids using the PEG/lithium acetate protocol. This *CDC14-mTFP1/WHI5-mKok/TUP1-mNeptune2.5/ mNeonGreen-ATG8* strain had WT kinetics for proliferation and sporulation.

Strains harboring Atg13-6SA, Atg13-2SA, or Atg13-PxL mutants were made by introducing mutated pRS303-harbored Atg13 or HA(or mScarlet)-Atg13 into *HIS* locus of parent strains, in which endogenous *ATG13* coding sequence was deleted by PCR-mediated knock-out with *URA/NAT* marker cassette.

### Media

The following media were used in this study. YPD (2% peptone, 1% yeast extract, 2% glucose), YPA (2% peptone, 1% yeast extract, 2% potassium acetate), SD (0.67% yeast nitrogen base, 2% glucose, auxotrophic amino acids and vitamins), and standard sporulation medium SPM (0.6% potassium acetate, pH 8.5). The SD-dropout medium was made with dropout stock powder lacking histidine, tryptophan, leucine, uracil, and/or methionine. Yeast cell nitrogen starvation experiments were performed in a synthetic minimal medium lacking nitrogen, SD-N (0.17% yeast nitrogen base without amino acids, 2% glucose).

### Plasmid Construction

Plasmids used in this study are listed in Table S2.

pRS303-Atg13 was constructed by subcloning PCR amplified regions of genomic *ATG13* with its promoter and terminator into NotI/SalI restriction sites of pRS303. 3HA, mScarlet, or mCherry fragments were fused to the N-terminus of *ATG13* by Gibson assembly cloning. Mutagenesis of all six Serine sites and 3 PxL sites, as shown in Table S2, were introduced by overlap extension (OE) PCR. pRS303-mScarlet-Atg8 was constructed by replacing the 2xGFP region in pRS303-2xGFP-Atg8 with mScarlet via Gibson assembly cloning.

Vectors for bacterial expression of 6His-3FLAG-tagged Cdc14 and GST-tagged Cdc14 were created by subcloning PCR amplified regions of the genomic Cdc14 encoding region into the NdeI/XhoI restriction sites of pET28a and BamHI/XhoI restriction sites of pGEX-6P-1, respectively. C283S mutation was introduced into pET28-Cdc14 by OE PCR.

Plasmids for C-terminal fluorophore tagging of Whi5-mKOk, Tup1-mNeptune2.5, and Cdc14-mTFP1 were confirmed by sequencing and restriction analysis and were originally reported in (Arguello-Miranda et al., 2018). pRS304 *ATG8p-mNeonGreen-ATG8-ATG8ter* was created by replacing the *Pac*I-2xyeGFP-*Bam*HI sequence in *pRS304-ATG8p-2xyeGFP-ATG8-ATG8ter* (Arguello-Miranda et al., 2021) with a *Pac*I-mNeonGreen*-Bam*HI sequence amplified from pYLB10 (Arguello-Miranda et al., 2018). The pRS423-Spo74-GFP was constructed by subcloning the Spo74-GFP fragment of pRS425-Spo74-GFP, which was a gift from Soni Lacefield in Indiana University (5), into SalI/SacI restriction sites of pRS423.

### Protein Expression and Purification

#### Recombinant protein purification

pET-based protein expression in BL21 (DE3) *E. coli* cells was induced by IPTG as described previously (Wang et al., 2010). 6His-3Flag-Cdc14 and 6His-3Flag-Cdc14m were expressed at 16°C overnight. Cells were collected by centrifugation, resuspended in bacteria lysate buffer (BLB: 20 mM Tris-HCl pH 8.0, 500 mM NaCl, 20 mM Imidazole, 10% glycerol, 5 mM 2-ME, 1 mM PMSF), and lysed using a High-Pressure Cell Press Homogenizer (Avestin Emulsiflex-C5). The lysate was supplemented with 0.1 mg/ml DNase I, 1 mM MgCl_2_, and 0.1% Triton-X 100 and cleared by centrifugation at 10,000g for 10 min at 4°C. The supernatant was applied to an NTA-Ni column (Qiagen, 30230), and washed three times with BLB. The 6xHis-proteins were eluted with 20mM Tris-HCl pH 8.0, 500 mM NaCl, 300 mM Imidazole, 10% glycerol and 5 mM 2-ME. The GST-Cdc14 constructs were expressed at 16°C overnight and the protein purification followed the same procedure as for 6His-3Flag-Cdc14 using glutathione-Sepharose beads (Millipore-Sigma, 17075601) and the following buffers: Lysate buffer for GST-Cdc14 (GSTLB) contained 20mM Tris-HCl pH 8.0, 500 mM NaCl, 10% glycerol, 5mM 2-ME, 1mM PMSF. Elution buffer contained GSTLB + 50 mM glutathione. The eluted proteins were then separated by Superdex 200 Increase 10/300 GL column (GE Healthcare) that was equilibrated in size exclusive buffer (SEC: 50 mM Tris/HCl pH 7.5, 500 mM NaCl, 10% glycerol, 5mM 2-ME). Finally, purified proteins were concentrated in SEC and frozen in liquid nitrogen.

#### IP of proteins expressed in yeast

Cells grown in SPM or YPD medium were pelleted at 3,000 × g for 5 min, 4°C, washed with ice-cold distilled water containing 1mM PMSF. The pellets were resuspended in ice-cold yeast lysis buffer (YLB: 50mM HEPES-KOH, pH 6.8, 150 mM KOAc, 2 mM MgCl_2_, 1 mM CaCl_2_, 0.2 M sorbitol, 10mM PMSF, 2x protease inhibitor cocktails [Roche, 11873580001]), dropped into liquid nitrogen, ground using a Retsch ball mill (PM100 or Retsch MM400) and stored at −80 °C for immunoprecipitation (IP), as well as for atg1 kinase assay and immunoblotting (IB).

To isolate the FLAG-Atg1-as complex, Frozen cell lysate powder (∼ 500 OD600 units) was thawed in 2ml 1 × IP buffer (50 mM HEPES-KOH (pH 6.8), 150 mM K_2_OAc, 2 mM MgOAc, 1 mM CaCl_2_, 15% glycerol, 1% NP-40, 1x protease inhibitor cocktail, 1x phosphatase inhibitors) and cleared twice by centrifugation at 1,000 × g for 5 min at 4 °C. 10 µl of Protein G Dynabeads (Invitrogen) that were loaded with mouse anti-FLAG M2 antibody (Sigma, F3165) were then added to the cell extract, incubated for 3 h at 4 °C with constant agitation. The beads were collected, washed 5 times with IP buffer containing 1% NP-40 and 1 × phosphatase inhibitors, and bound proteins were eluted with 25 µl 1 mg/ml 3 × FLAG peptide (Sigma, F4799) in IP buffer containing 1% NP-40 and 1 × protease inhibitors at 4 °C. Eluates were aliquoted, frozen in liquid nitrogen, and stored at −80 °C. The purified FLAG-Atg1 complex was resolved by SDS-PAGE and analyzed by Sypro Ruby staining (Thermo Fisher, S12000).

To purify 3HA-Atg13 and its variants, 0.4g frozen cell lysate from cells grown in SD medium (Fig. 4C,4E, 4F) or SPM (Fig. 4D) was thawed on ice and supplemented with 0.8ml IP buffer. Extracts were cleared twice by centrifugation at 1,000x g for 5 minutes at 4 °C. Supernatants were then incubated with 5 µl protein G magnetic beads with bound anti-HA antibody, for 3 hours at 4 °C with constant agitation. After 5 times washing by IP buffer, resins were resuspended in 100 µl of IP buffer and used for: (1) phosphatase treatment (Fig. 4E,4F); (2) Cdc14 pull-down (Fig. 4C); (3) Immunoblotting (IB) (Fig. 4D).

### Sporulation, meiotic division synchronization, and vegetative growth

A single colony of yeast strains from the YPD plate was picked, spread on YPG (3% glycerol) plates, and grown at 30 °C for 2 days. Colonies grown out were spread to YPD plates and grown until cells formed a lawn (∼ 24 hours). Cells on plates were collected and suspended in YPA liquid medium (OD600=0.3) and grown for 14-16h at 30 °C. Cells were then pelleted, washed twice with water, and resuspended in SPM to final OD600= 3. For meiotic division synchronization, following incubation in SPM for 12 hours, strains containing *GAL-NDT80 GAL4-ER* were released from prophase I arrest by addition of 1μM β-estradiol to induce Ndt80 expression. To assess the percentage of spore formation, cells with matured spores were counted after 48 h in SPM at room temperature. At least 300 cells of each strain were counted by bright field microscopy (Olympus BX40, 40x objective).

For single-cell time-lapse imaging, the OAM835 strain was inoculated to 0.03 OD600 and grown for 23 hours or until the OD600 reached 2.5. 70 µl of the culture were sonicated (4-6 s at 3 W) and transferred to a Y04C CellASIC microfluidic device (http://www.cellasic.com/) set to 25 ^°^C. Two pulses of cells were loaded (8 psi for 5 s) and then exposed to a meiosis-inducing medium (0.6% potassium acetate, delivered at 0.6 psi flow rate, pH 8.0 adjusted with 0.125 M Na_2_CO_3_ immediately before the experiment) for 24 h.

For vegetative growth experiments, cells were cultured in YPD or SD medium to log phase (OD=0.6∼1.0). To induce expression of pZD promoter-controlled genes (Cdc14 variants, Atg13 8SA, GFP-Atg8, and mCherry-GFP), 1 μM β-estradiol (10 mM stock in ethanol, Sigma E2758) was added. To inhibit Atg1-as, 5 μM of 1-NM-PP1, was added (10 mM stock in DMSO, APExBIO, B1299), as needed.

### Immuno-blotting (IB) analysis

3 OD600 of yeast cells were collected by centrifugation, resuspended in 100µl 2x SDS sample buffer and boiled at 95°C for 5 min [2xSSB: 62.5 mM Tris-HCl, pH=6.8; 2% SDS; 0.05% BPB; 10% Glycerol; 5% 2-Mercaptoethanol; 1× Protease Inhibitor Cocktail (Roche, 11873580001), 1 mM PMSF]. Proteins were separated by SDS-PAGE (90 min at 150V) using 4-20% Criterion™ TGX Stain-Free™ gel (Bio-Rad, 5678095) and electro-blotted onto nitrocellulose membranes (Bio-Rad, 1620115) using a Trans-blot SD semi-dry transfer cell (Bio-Rad, 1703940). After blocking with 5% skim milk in TBST and incubated with respective primary antibodies followed by HRP or StarBright® B700 conjugated secondary antibodies. For detection and quantitative analysis, the IB images were captured by ChemiDoc™ MP imaging system (Bio-Rad, 12003154), and analyzed using Image Lab™ (Ver. 6.0.1) software (Bio-Rad). 7.5% mini TGX Stain-Free™ protein gels (Bio-Rad, 4561026) were used to separate phosphorylated and dephosphorylated bands of Spo74-GFP.

We used the following antibodies: Recombinant monoclonal anti-Thiophosphate ester rabbit IgG [51-8] (Abcam, ab92570), monoclonal anti-GFP mouse IgG (Roche, 11814460001, 1:5000), polyclonal anti-Hexokinase 1 rabbit IgG (United States Biological, 169073, 1:10000), polyclonal anti-NDT80 rabbit IgG (1:10000), monoclonal anti-HA mouse IgG1 (Thermo Fisher, 21683, 1:3000), monoclonal anti-FLAG M2 mouse IgG (Millipore-Sigma, F3165, 1:5000), polyclonal anti-FLAG rabbit IgG (Millipore-Sigma, SAB4301135, 1:5000), polyclonal anti-GST rabbit IgG (Millipore-Sigma, G7781), StarBright® B700 labeled goat anti-mouse IgG secondary antibody (Bio-Rad, 12004158, 1:5000), HRP conjugated goat anti-mouse IgG (Bio-Rad, STAR207P) and StarBright® B700 labeled goat anti-rabbit IgG (Bio-Rad, 12004161, 1:5000).

### *In vitro* Cdc14 phosphatase activity assay

500mM p-Nitrophenyl Phosphate (pNPP, NEB, P0757S) substrate solution was diluted to 20 mM in phosphatase reaction buffer (PRB: 20mM Tris pH7.5, 4 mM MgCl_2_, 1 mM EGTA, 0.02% 2-ME, 0.1 mg/ml BSA) to yield pNPP working stock. Recombinant 6His-3Flag-Cdc14/6His-3Flag-Cdc14m proteins at various concentrations (0.25-80 µg/ml prepared in PRB) were mixed with the pNPP working stock in 96-well plates to a final concentration of pNPP of 10 mM. The reaction mix was incubated for 5-45 min at room temperature and stopped by 1 N (final concentration) NaOH. Absorbance at 405 nm was measured on a BioTek Gen5 microplate reader. 0.4 µg/ml λ phosphatase (NewEngland Biolabs, P0753S) and 1x phosSTOP cocktail (Millipore-Sigma, 4906845001) were added to control reactions.

### *In vitro* dephosphorylation of 3HA-Atg13

20 µl of 3HA-Atg13 immobilized on protein G magnetic beads, was incubated with 50 µg/ml 6His-3Flag-Cdc14, 6His-3Flag-Cdc14m or 10 µg/ml λ PPase in reaction buffer (50 mM HEPES-KOH pH 6.8, 150 mM K_2_OAc, 350 mM NaCl, 2 mM MgOAc, 1 mM CaCl_2_, 15% glycerol, 0.1% Triton X-100) in 30 °C water bath for 20 min. The reaction was stopped by the addition of 1x phosSTOP cocktail and used for Atg1 kinase assay (Fig. 4F) or Flag-Atg1 pulling down assay (Fig. 4E).

### Atg1 pull-down by 3HA-Atg13

Frozen cell lysates from Flag-Atg1-as cells cultured in YPD to log phase (OD=0.6∼1.0) were thawed, diluted with an equal volume of IP buffer (150 mM HEPES-KOH pH 6.8, 150 mM K_2_OAc, 2 mM MgOAc, 1 mM CaCl_2_, 15% glycerol, 1% NP-40, 1x Roche protease inhibitor cocktail and 1 x phosSTOP cocktail) and cleared twice by centrifugation at 1,000x g for 5 min at 4 °C. 20 µl of magnetic beads coated with 3HA-Atg13 were incubated with 100 µl Flag-Atg1-as cell extracts for 4 hours at 4 °C. Resins were collected and washed 3 times with ice-cold IP buffer. Proteins were eluted by boiling in SSB, separated by SDS-PAGE, and analyzed by IB using appropriate antibodies.

### Cdc14/Cdc14m pull-down by 3HA-Atg13

20 µl of the bead-immobilized 3HA-Atg13 were incubated with 50 µg/ml 6His-3Flag-Cdc14 or 6His-3Flag-Cdc14m in 50 ul binding buffer (50 mM HEPES-KOH pH 6.8, 150 mM K_2_OAc, 350 mM NaCl, 2 mM MgOAc, 1 mM CaCl_2_, 15% glycerol, 0.1% Triton X-100, 0.05% 2-ME, 1x Roche protease inhibitors, 1x PhosSTOP cocktail) overnight at 4 °C. The beads were washed 3 times with binding buffer, boiled in SSB to elute bead-bound proteins that were analyzed by SDS-PAGE and IB with appropriate antibodies.

### Assay of Atg1-as kinase activity

#### In cell lysates (Fig. 2B, 2E, 3E)

Cell lysates were mixed with 1 × kinase buffer (150 mM K_2_OAc, 10 mM MgOAc, 0.5 mM EGTA, 5 mM NaCl, 20 mM HEPES-KOH pH 7.3, 5% glycerol) in equal volume (wt/vol) and thawed on ice. After two centrifugations at 1,000x g for 5min at 4 °C, supernatants were mixed with equal volumes of 2 × kinase mix (kinase buffer, energy mix [90 mM creatine phosphate, 2.2 mM ATP, 0.45 mg/ml creatine kinase] and 0.2 mM N6-phenylethyl-ATPγS [N6-PhEt-ATPγS, BLG-P026-05]) and incubated for 1.5 h at room temperature. Reactions were quenched with 20mM EDTA and then alkylated with 2.5 mM paranitrobenzyl mesylate (PNBM, Abcam) for 45 min at room temperature, boiled in SSB, and analyzed by immunoblotting. Thiophosphorylated substrates were identified by immunoblotting with a rabbit anti-thiophosphate ester primary antibody [51-8] (Abcam, ab92570) and StarBright® B700 labeled goat anti-Rabbit IgG secondary antibodies.

#### In cell lysates + 3HA-Atg13 (Fig. 4F)

Following bead-immobilization and phosphatase treatment as described above, 5 µl of 3HA-Atg13 on magnetic beads was incubated with 10 µl GFP-Atg1-as cell extracts in 1 × kinase buffer for 15 min at room temperature with constant agitation, followed by atg1-as kinase assay.

#### In purified Flag-Atg1-as complex (Fig. 4A)

25 µg/ml or 50 µg/ml recombinant Flag-Cdc14 were incubated with 2 µl Flag-Atg1-as complex, supplemented with 1 × kinase buffer to 15 µl, for 20 min in a 30 °C water bath, followed by atg1-as kinase assay as described. 1x phosSTOP cocktail was added to inhibit Cdc14 phosphatase as control.

### Spo74-GFP immunoblotting, immunoprecipitation, and *in vitro* dephosphorylation

100 OD Cells harboring pSPO74-Spo74-GFP were harvested at indicated time points (Fig. 6) in SPM. After ball milling, frozen cell powder was dissolved in 2x SSB, heated at 95 °C for 5 minutes, and analyzed by SDS-PAGE and immunoblotting. For immunoprecipitation, cell lysates were suspended in 2x volume (vol/wt) of IP buffer and cleared twice by centrifugation at 1,000 x g, 5 minutes at 4 °C. Supernatants were incubated with anti-GFP antibodies bound to protein G magnetic beads for 3 hours at 4 °C and washed 3 times with 1x IP buffer. Resins were incubated with 20 µg/ml GST-Cdc14 (+/-phosSTOP cocktail) or 10 µg/ml λ PPase at 30 °C in a water bath for 30 minutes. Reactions were terminated by heating samples in 2x SSB.

### Fluorescence microscopy

Tubulin immunofluorescence analysis as described earlier (Wang et al., 2020). In brief, after Ndt80 was induced, cells were fixed every 30 minutes by cold 3.7% formaldehyde in PBS. Fixed cells were washed with 0.1 M KPi (4.84 g/L potassium phosphate dibasic; 9.83 g/L potassium phosphate monobasic; pH 6.4) buffer and 1.2 M sorbitol citrate solution, and then digested by 1.4 mg/ml 20T zymolyase (amsbio, 120491) and glusulase (PerkinElmer, NEE154001EA). Cells were spread on poly-L-lysine (Sigma, P8920) coated glass slides and incubated with anti-tubulin Alexa Fluor® 594 antibody (Abcam, dilution 1:200) for 2 hours, and mounted in ProLong Gold antifade reagent containing DAPI (ThermoFisher, P36935). Spindle morphologies were then classified by the following morphological features: metaphase I cells was defined as cells with a single DAPI mass spanned by thick, short, and bipolar meiotic spindle microtubules (approximately 2–3μm in length). Anaphase I cells was defined as cells with two parts of distinct (though not always separated) DAPI masses, and a single long spindle that spans both DAPI masses. Metaphase II cells were defined as cells with two separate DAPI masses with each spanned by a bipolar, thick and short meiotic spindle. Anaphase II cells were defined as cells with four distinct (though not always separated) parts of DAPI masses with two long spindles (usually crossed). Images were acquired using a Zeiss Axio Imager Z2 microscope equipped with a Photometrics CoolSnap HQ2 CCD camera, a Zeiss PLAN APOCHROMAT 63×/0.9 NA oil immersion objective, and filters for DAPI and Alexa594. At least 100 cells were counted at each time point, to determine the percentage of cells at each meiotic cell stage.

For live-cell imaging, 10 µl of cells growing in SD medium or SPM were spread on a slide and imaged immediately using a Zeiss Axio Imager Z2 microscope equipped with a Photometrics CoolSnap HQ2 CCD camera, a Zeiss PLAN APOCHROMAT 63×/0.9 NA oil immersion objective and filters for GFP/FITC, mCherry/Alexa594. Strains harboring Cdc14-GFP(*IN*)/mScarlet-Atg13 and all strains with Spo74-mTFP were imaged using 3i spinning disk confocal microscope, using a 63x oil immersion objective with mTFP filter (473-488/10), mNeonGreen filter (515-542/27), mRuby filter (561-617/73) and GFP filter (413-525/80). For imaging of cells expressing Spo74-mTFP, 5 µM 1NM-PP1 or 200 µM Chloroquine (Millipore-Sigma, C6628) were added 3 hours after Ndt80 induction, if needed. Z-stack (5-10 planes, 0.5 µm/plane) images were acquired if needed. Images were processed using ImageJ except for the analysis of single-cell images as described above.

### Time-Lapse Microscopy, image processing and quantification of cellular features

Time-lapse images of microfluidics experiments were acquired using a motorized ZEN software-operated Zeiss Observer Z1 microscope with temperature control, a Definite Focus 2.0 system, 40X Zeiss EC Plan-Neofluar 1.3 NA oil immersion objective, and an AxioCam HR Rev 3 camera. At least 3 fields of five replicates were imaged. A set of custom dichroic mirrors and bandpass filters were used for fluorescence imaging with exposure times of mTFP1 125 ms, mNeonGreen 175 ms, mKO*κ* 200 ms, mNeptune2.5 75 ms, and phase contrast 40 ms. LED light sources were kept at 25% intensity except for mNeptune2.5 at 50%.

ZEN pro software (Zeiss) was used to collect 2 x 2 binning non-compressed TIFF format images that were converted to double format before feature extraction. A 3 x 3 structured *medfilt2()* filter removed shot noise. Cells were segmented with a previously described pipeline (Doncic et al., 2013, Wood and Doncic, 2019), background quantification was defined as the median intensity of the space not occupied by cells. A 2D Gaussian fit to the brightest pixel of nuclear proteins (Tup1 or Whi5) was used to calculate nuclear parameters (Doncic et al., 2013). Nucleus Cdc14 released was computed by a 2D gaussian filtered-based algorithm that labels the nucleus’ position using the brightest nuclear pixel associated with the Cdc14 signal. The time of Cdc14 release was defined by this algorithm as the time point where the fluorescence intensity of the nuclear Cdc14 signal irreversibly decreases by 20%. mNeonGreen foci were detected with a two-step algorithm: First, the vacuole was segmented using a label-free intensity threshold-based algorithm directly from gaussian-blurred phase contrast images. Second, a dynamic intensity-based 2D gaussian filter algorithm was used to quantify the number, size (in pixels), and other properties of the algorithmically defined Atg8 puncta. Foci were defined as vacuolar or non-vacuolar if they overlapped with the area algorithmically identified as the vacuole. The code for label-free vacuolar tracking and extraction and mNeonGreen-Atg8 foci quantification is available at: https://github.com/alejandrolvido/Vacuole_buster

